# Cryo-EM structures of a synthetic antibody against 22 kDa claudin-4 reveal its complex with *Clostridium perfringens* enterotoxin

**DOI:** 10.1101/2023.06.12.544689

**Authors:** Satchal K. Erramilli, Pawel K. Dominik, Chinemerem P. Ogbu, Anthony A. Kossiakoff, Alex J. Vecchio

## Abstract

Claudins are a family of ∼25 kDa membrane proteins that integrate into tight junctions to form molecular barriers at the paracellular spaces between endothelial and epithelial cells. Humans have 27 subtypes, which homo- and hetero-oligomerize to impart distinct properties and physiological functions to tissues and organs. As the structural and functional backbone of tight junctions, claudins are attractive targets for therapeutics capable of modulating tissue permeability to deliver drugs or treat disease. However, structures of claudins are limited due to their small sizes and physicochemical properties—these traits also make therapy development a challenge. We have developed a synthetic antibody fragment (sFab) that binds human claudin-4 and used it to resolve structures of its complex with *Clostridium perfringens* enterotoxin (CpE) using cryogenic electron microscopy (cryo-EM). The resolution of the structures reveals the architectures of 22 kDa claudin-4, the 14 kDa C-terminal domain of CpE, and the mechanism by which this sFab binds claudins. Further, we elucidate the biochemical and biophysical bases of sFab binding and demonstrate that this molecule exhibits subtype-selectivity by assaying homologous claudins. Our results provide a framework for developing sFabs against hard-to-target claudins and establishes the utility of sFabs as fiducial markers for determining cryo-EM structures of this small membrane protein family at resolutions that surpass X-ray crystallography. Taken together, this work highlights the ability of sFabs to elucidate claudin structure and function and posits their potential as therapeutics for modulating tight junctions by targeting specific claudin subtypes.

## Introduction

The claudin family of integral membranes proteins constitute the structural and functional backbone of endothelial and epithelial tight junctions in vertebrates. In mammals, 24+ claudin subtypes self-assemble, forming large homo- and/or heteromeric complexes that create barriers to the transport of small molecules and ions through intercellular spaces. Claudins have four transmembrane helices (TMs), which, along with two extracellular segments (ECS), enable tight junction assembly by simultaneously interacting laterally within the same membrane (cis) and across intercellular space with claudins on adjacent cell membranes (trans) [1, 2]. Cis and trans interactions between the various sized and shaped claudin subtypes govern the frequency and morphology of tight junction strands, specifically tuning small molecule permeability to impart distinct molecular properties and physiological functions to tissues and organs [3]. Altering tight junction barrier permeability via transient disruption of claudin/claudin interactions is being investigated as a strategy to treat tight junction-linked diseases or to deliver drugs through these highly-restrictive impediments, like through the blood-brain barrier [3–5]. Targeting specific claudins with therapeutic agents, which would be required for tissue- or disease-specific treatments, however, remains a challenge.

Claudin-4 functions in normal physiological processes largely through its role in increasing the complexity of tight junctions and tuning the paracellular permselectivities of various tissues [6]. In lung epithelium, it regulates the paracellular barrier to modulate fluid clearance [7]. In kidney epithelium, it forms a chloride-selective channel vital for chloride reabsorption via interactions with claudin-8 [8]. Claudin-4 in intestinal epithelium assists M cells, specialized cells that capture microbial microparticles and transport them through epithelium to immune cells, by acting as a structural component for M cell transcytosis vesicles [9]. Dysregulated expression of claudin-4 has been detected in bladder, breast, colorectal, gastric, lung, ovarian, pancreatic, prostate, and thyroid cancers [10, 11]. Claudin-4 functions diversely in normal versus disease states, making it a primary target of therapeutics to modify claudin-4 structure and function.

In the gastrointestinal tracts of mammals, claudin-4 is also the endogenous receptor of *Clostridium perfringens* enterotoxin (CpE) [12, 13]. CpE infection causes a highly prevalent form of food poisoning by disrupting tight junctions in the gut [14–16]. Claudin structural and sequential homology enables other subtypes to also bind CpE with high-affinity [17, 18]. We have previously pinpointed a conserved sequence that spans claudin 3D structure called the “cCpE-binding motif”, which imparts claudin selectivity through its recognition by CpE’s C-terminal domain (cCpE) [19]. The N-terminal domain of CpE functions in cytotoxicity by structurally rearranging into a membrane-spanning β-barrel that induces cytotoxicity by creating a calcium-selective pore [15]. Although cCpE has no cytotoxic domain and does not kill cells, it is not harmless. Delivery of cCpE to the basolateral compartment of epithelial monolayers sequesters claudin-4 away from tight junctions, disrupting its barrier function [20]. The natural abilities of CpE and cCpE to bind claudins in a subtype-selective and high-affinity manner, while disrupting claudin assemblies and thus the barrier function of tight junctions, make these enterotoxins prospective modulating agents of tight junctions containing claudins with receptor capacities, like claudin-4. Specifically, enterotoxins are being used to target claudin-4 in ovarian, pancreatic, and other tissue-specific cancers [11, 21–28]. Modified enterotoxins have been employed to target claudins with low receptor capacities like claudin-5, to modulate opening of the blood-brain barrier [29–33]. Claudin targeting by novel anti-claudin binders or by CpE enterotoxins thus represents a strategic and promising approach for the development of imaging probes, drugs, and therapeutics to treat tight junction-linked diseases.

Advancement of claudin binders is slow due to their small size, minute extracellular masses, physicochemical properties, homologous architectures, and dearth of experimentally derived structures—which could facilitate therapeutic efforts. To date, structures of two human and three murine claudins have been experimentally determined using X-ray crystallography to resolutions that illuminate their biology (<4.0 Å) [34–39]. The features that make claudins challenging to target with therapeutics also make them recalcitrant to structural determination—especially their proportionally low soluble ECS mass-to-hydrophobic TM ratios. Of the 27 human claudins, which range in size from 22-34 kDa, claudin-4 is the second smallest with 209 amino acids that amass to 22,077 Da. With over 50% of claudin-4s mass being buried within membranes or disordered as termini, there is little “targetable” tertiary structure presented as antigenic or binding surfaces for drugs. This small area is also all that is available to form the requisite protein/protein interactions for crystal nucleation. Detergents used to solubilize claudins for *in vitro* and structural studies additionally block protein packing, crystallization, and drug-like molecules from binding claudins effectively. Developing approaches and/or molecules capable of circumventing the low mass-to-membrane ratios of claudins thus holds promise to augment both structural biology and tight junction modulating therapeutic efforts.

We have developed a platform to coordinate structure determination and molecular targeting of claudins using synthetic antibody fragments (sFabs). Applying a sFab-encoding phage display library against detergent soluble human claudin-4 (hsCLDN-4) bound to cCpE, we have discovered three sFabs termed CpE Obstructing Proteins (COPs). We previously have shown that COP-2 and COP-3 bind cCpE, but not hsCLDN-4, and used these COPs to determine low resolution (5-7 Å) structures of hsCLDN-4/cCpE complexes using single-particle cryogenic electron microscopy (cryo-EM) at 200 kV [40]. Here, we describe COP-1, a hsCLDN-4-selective sFab, and employ it to reveal structures of hsCLDN-4/cCpE complexes using cryo-EM at 300 kV to 2.6 Å—a resolution that exceeds all claudin/cCpE structures determined previously with X-ray crystallography. Structural, biophysical, and biochemical analyses aid elucidation of the claudin/cCpE interactions required for CpE-induced cytotoxicity and COP-1s distinct hsCLDN-4-selective binding mechanism. Notably, COP-1 identifies hsCLDN-4 by penetrating the hydrophobic environment to bind TM-proximal regions. COP-1 thus naturally circumvents the membrane barrier, exposing new antigenic regions of claudins previously considered inaccessible. COP-1s unique binding capability can be used as a prototype to bolster drug development efforts to advance new therapies that modulate tight junctions in tissue-specific ways through claudin-selective targeting.

## Results

### Development and Sequence of COP-1

COP-1 was co-discovered with two anti-cCpE sFabs, COP-2 and COP-3, using a phage display library encoding sFabs with variable complementarity-determining regions (CDRs) targeted against hsCLDN-4 solubilized in n-dodecyl-β-D-maltopyranoside (DDM) in complex with cCpE [40, 41]. COP-1 was sequenced along with COP-2 and -3. Sequence alignments revealed the expected variability in the CDR regions of the light (L) and heavy (H) chains (***SI Appendix*, Fig. S1**). Sequence conservation is high in CDR-1 and CDR-2 of both L and H chains, with only minor alterations from Ser to Tyr. However, the CDR-3 regions in both chains of COPs vary significantly. The CDR-L3 of COP-1 has one aromatic residue compared to three and five in COP-2 and -3, and is truncated by one and two amino acids, respectively. COP-1s CDR-H3 has nine aromatic residues compared to seven and six in COP-2 and -3; there is also a one amino acid insert in COP-1, a Pro. The Pro precedes four sequential Trp residues. At this point, COP-1 was expressed and purified to characterize its ability to bind various antigens *in vitro*. When purifying COP-1, we found it was not as soluble as COP-2 and -3 and benefits from adding detergents (data not shown). The string of Trp residues and the increased number of hydrophobic aromatic side chains in CDR-H3 may explain this behavior and direct COP-1s binding function.

### Biochemical Analysis of COP-1

Once isolated, we biochemically characterized COP-1s ability to bind hsCLDN-4 and hsCLDN-4/cCpE complexes *in vitro* using size-exclusion chromatography (SEC) in DDM and DDM/cholesteryl hemisuccinate (CHS). The presence of CHS had no effect on binding results apart from earlier elution times of complexes due to increased mass caused by the addition of CHS to DDM micelles. This study showed that cCpE binds hsCLDN-4; COP-1 binds well to hsCLDN-4/cCpE complexes but only slightly to hsCLDN-4 alone; and that COP-1 and COP-2 bind hsCLDN-4/cCpE complexes simultaneously (**Fig. 1A** and **B**). We had previously shown that COP-2 binds only to cCpE, which suggested that COP-1s epitope was unique and that it may not bind cCpE [40]. The data further showed that when hsCLDN-4/cCpE/COP-1 were complexed, excess COP-1 (peak 6) and cCpE (peak 7) did not result in a peak at the elution volume of cCpE/COP-2 complexes (peak 5), which indicated that COP-1 did not bind cCpE alone unlike COP-2 (**Fig. 1A**). These analytical studies provided a foundation to probe COP-1 binding further using structural and biophysical techniques.

**Figure 1.**
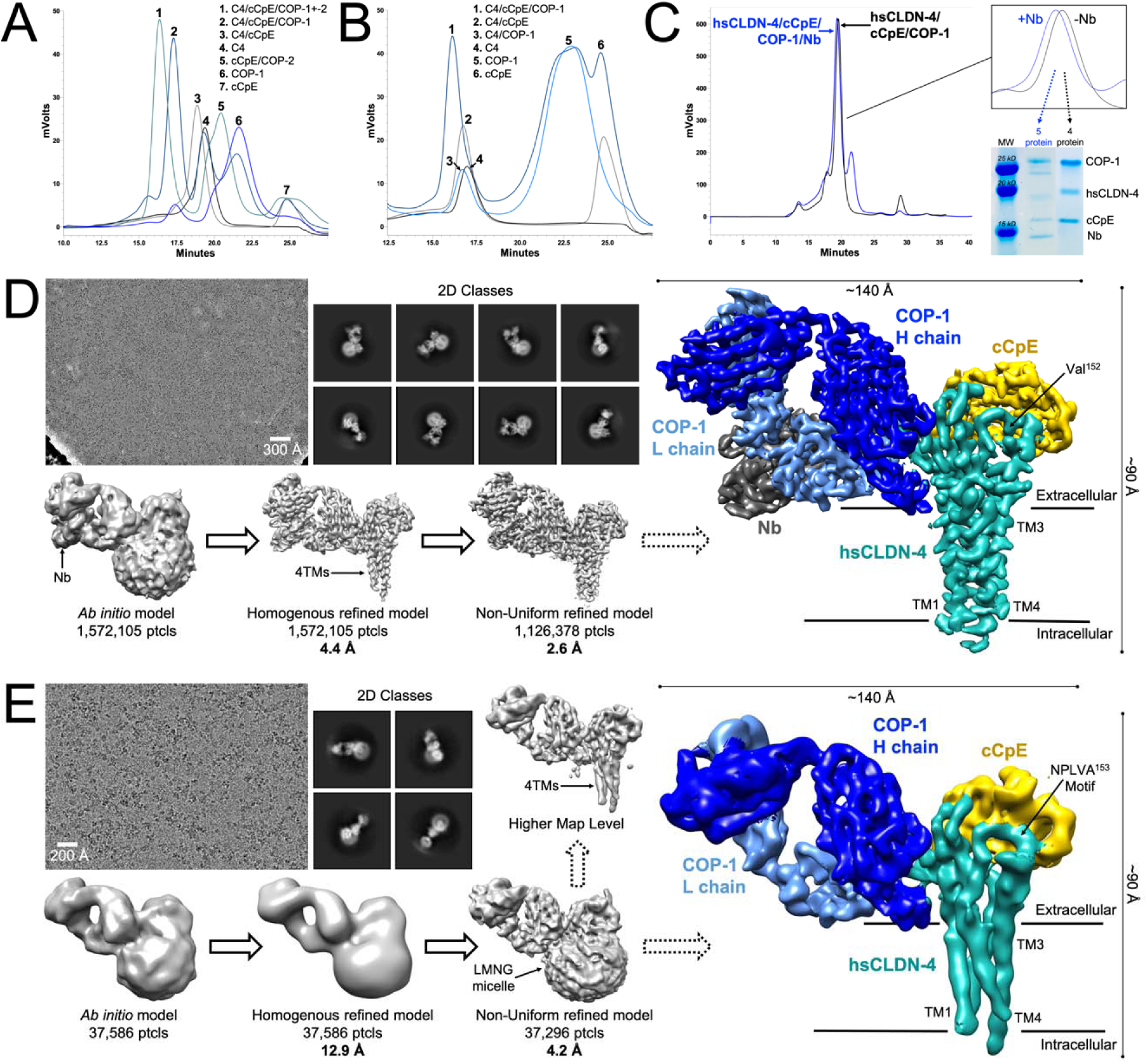
Biochemical Characterization and Preparation, and Structure Determination of Claudin-4/cCpE/COP-1 Complexes. (A) Analytical size-exclusion (SEC) chromatograms depicting elution profiles and peak retention times of various proteins and complexes between hsCLDN-4, cCpE, COP-1, and COP-2 in a mobile phase with DDM. (B) Analytical SEC chromatograms shown as in A but with a mobile phase containing DDM/CHS. Note that elution times of complexes in DDM/CHS are altered compared to A but that cCpE and COP-1 retention times are not. (C) Prep-scale SEC chromatograms of protein complexes used for cryo-EM with corresponding purity of the four- or five-protein complexes analyzed using SDS-PAGE. (D) Overview of cryo-EM analysis of the hsCLDN-4/cCpE/COP-1/Nb complex showing motion-corrected micrograph, 2D classes, and initial and final 3D reconstructions. The final map is shown in relation to a model membrane boundary and the five proteins are colored accordingly: hsCLDN-4 (teal), cCpE (gold), COP-1 H (blue) and L (light blue) chains, and Nb (grey). Dimensions of the complex in 2D are also provided. (E) Overview of cryo-EM analysis of the hsCLDN-4/cCpE/COP-1 complex as shown in D with final map colored identically but without Nb.

### Structure Determination of hsCLDN-4/cCpE/COP-1 Complexes by Single-Particle Cryo-EM

Because complexes of hsCLDN-4, cCpE, and COP-1 were preserved through SEC we sought to determine the basis of COP-1 binding and to discern whether sFabs could yield high resolution structures of claudins with single-particle cryo-EM, which we postulated previously [40]. If so, cryo-EM could prove or disprove whether interactions hypothesized to be vital for hsCLDN-4 and cCpE association from ∼3.5 Å crystal structures were present in an environment free from crystal-induced packing interactions [36, 39]. Cryo-EM, therefore, had potential to provide new insights into claudin biology and expedite structural biology workflows.

HsCLDN-4 was purified in 2,2-didecylpropane-1,3-bis-β-D-maltopyranoside (LMNG) detergent and bound to cCpE and COP-1 sequentially. This ∼87 kDa hsCLDN-4/cCpE/COP-1 complex was then bound to a ∼15 kDa anti-sFab nanobody (Nb) known to bind to the hinge region between the variable and constant domains of sFab L chains [42]. We surmised that the Nb may help by adding mass to yield a >100 kDa complex, giving a unique shape to hsCLDN-4/cCpE/COP-1 particles that could be readily distinguished in 2D classification, and decreasing flexibility inherent to sFab constant domains. Both hsCLDN-4/cCpE/COP-1 and hsCLDN-4/cCpE/COP-1/Nb were polished using SEC and eluted as four- or five-protein complexes (**Fig. 1C**). These complexes were applied to EM grids, vitrified, and subjected to structural analyses using single-particle cryo-EM.

We collected a dataset containing 5,039 movies of the hsCLDN-4/cCpE/COP-1/Nb complex first. During data processing, COP-1/Nb yielded identifiable particles in 2D classification and 3D reconstructions (**Fig. 1D**). Further processing generated an initial 4.4 Å map. This map showed that all five proteins were present and was of sufficient quality to visualize their secondary structural elements and corresponding side chains. Contouring this map revealed that density for the LMNG micelle gave way to the four TM helix bundle intrinsic to the claudin fold. Additional processing and refinement improved the map, which was finally resolved to a global resolution of 2.6 Å. All five proteins and hsCLDN-4s four TMs were visualized (**Fig. 1D**). Before data processing of the hsCLDN-4/cCpE/COP-1/Nb complex was complete we knew it had high resolution potential due to the quality of 2D classes. For this reason, we collected only a 1,071-movie dataset of the hsCLDN-4/cCpE/COP-1 complex. Processing resulted in 37,296 particles, which were refined to generate a map with a final resolution of 4.2 Å (**Fig. 1E**). The resolution of the hsCLDN-4/cCpE/COP-1 map was sufficient to resolve the hole between COP-1 domains, secondary structures of all four proteins, and the four TMs and NPLVA^153^ motif of hsCLDN-4 (**Fig. 1E**).

The cryo-EM maps from both complexes appeared similar except for the additional density that constitutes the Nb. This showed that the Nb had no influence on COP-1s binding pose to hsCLDN-4/cCpE. We built initial structural models into both cryo-EM maps starting from density corresponding to the hsCLDN-4/cCpE complex using PDB ID 7KP4, then added COP-1 or COP-1/Nb manually. These structures were then optimized to best fit the experimental maps using real-space refinement programs, constructing the final models of the hsCLDN-4/cCpE/COP-1 and hsCLDN-4/cCpE/COP-1/Nb complexes (see **Materials** and **Methods**). Because the Nb did not alter COP-1 binding and this map was 1.5 Å higher resolution, all subsequent structural analysis focused exclusively on the 2.6 Å hsCLDN-4/cCpE/COP-1/Nb complex.

### The High Resolution Cryo-EM Map Allows Visualization of 22 kDa Claudin-4

The utility of the 2.6 Å cryo-EM map became apparent immediately as we could visualize the complete tertiary structure of full-length hsCLDN-4 apart for its unstructured intracellular termini—180 of 209 residues (**Fig. 2**). This was significant because no experimentally determined claudin map had resolved all structural elements from TM1 to TM4, including a 2.4 Å crystal structure of mouse claudin-15 [34]. This map clearly resolved all four TMs, both ECS including the connecting loops, and showed the positions of 180 side chains (**Fig. 2** and ***SI Appendix*, Fig. S2A**). We performed local resolution analysis of the 2.6 Å map to approximate the resolution of different regions (**Fig. 3A**). This analysis showed that the average resolution of the map around hsCLDN-4 was <3.0 Å, with the intracellular loop (ICL) connecting TM2 to TM3 exhibiting the most dynamics. Local resolutions of other regions showed that the variable domain of COP-1 and its antigen binding surfaces were resolved to <2.4 Å while the rest of the complex was resolved to between 2.6-3.0 Å (**Fig. 3A** and ***SI Appendix*, Fig. S2B**). The map was thus sufficient for precise placement of the entire hsCLDN-4/cCpE/COP-1/Nb complex within its density, which allowed for structure elucidation of cCpE binding to hsCLDN-4 and of COP-1s binding mode (**Fig. 3B**).

**Figure 2.**
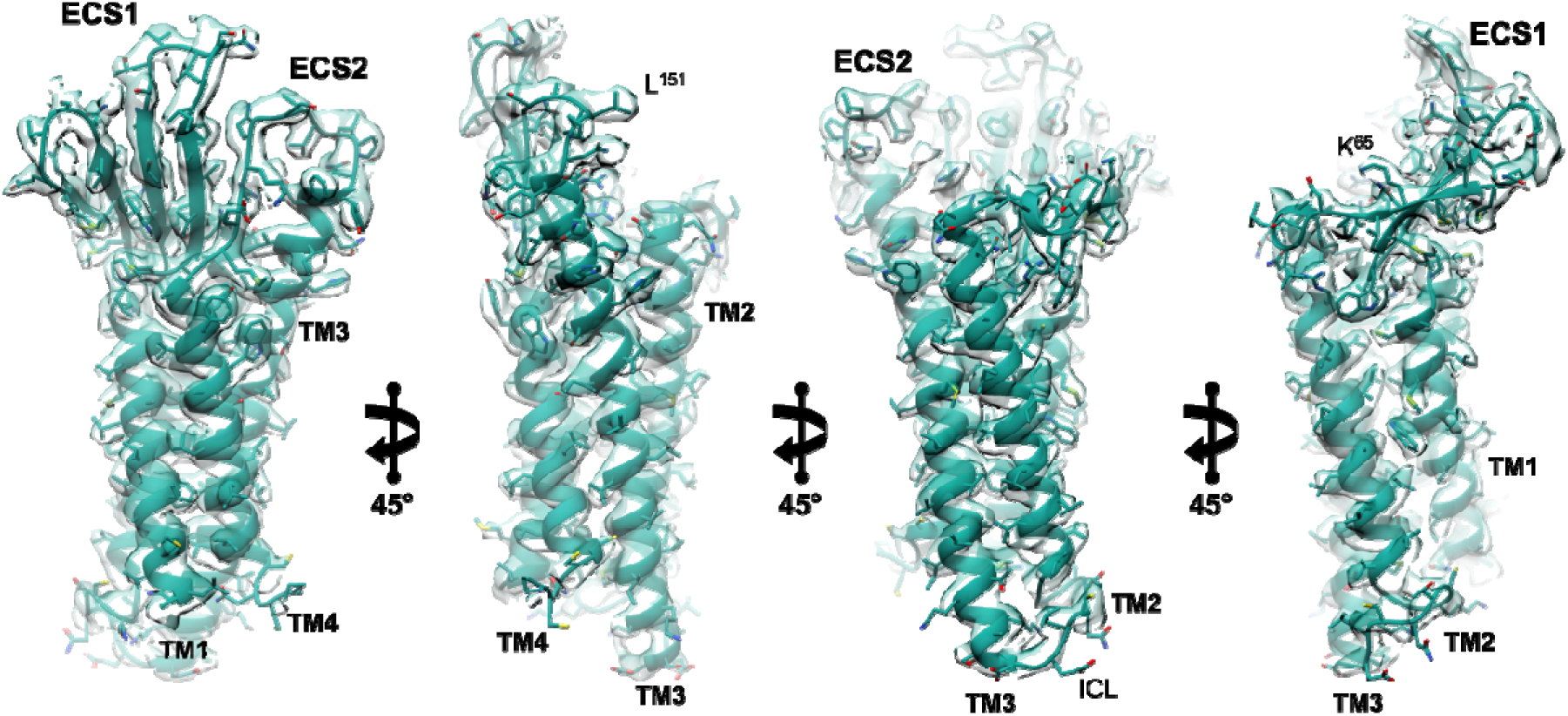
Visualization of the 22 kDa Claudin-4 Fold. The 2.6 Å cryo-EM map from the hsCLDN-4/cCpE/COP-1/Nb complex (teal) overlaid on top of the structural model of hsCLDN-4 (teal). Carbons (teal), oxygens (red), nitrogens (blue) and sulfurs (yellow) are colored accordingly on side chains.

**Figure 3.**
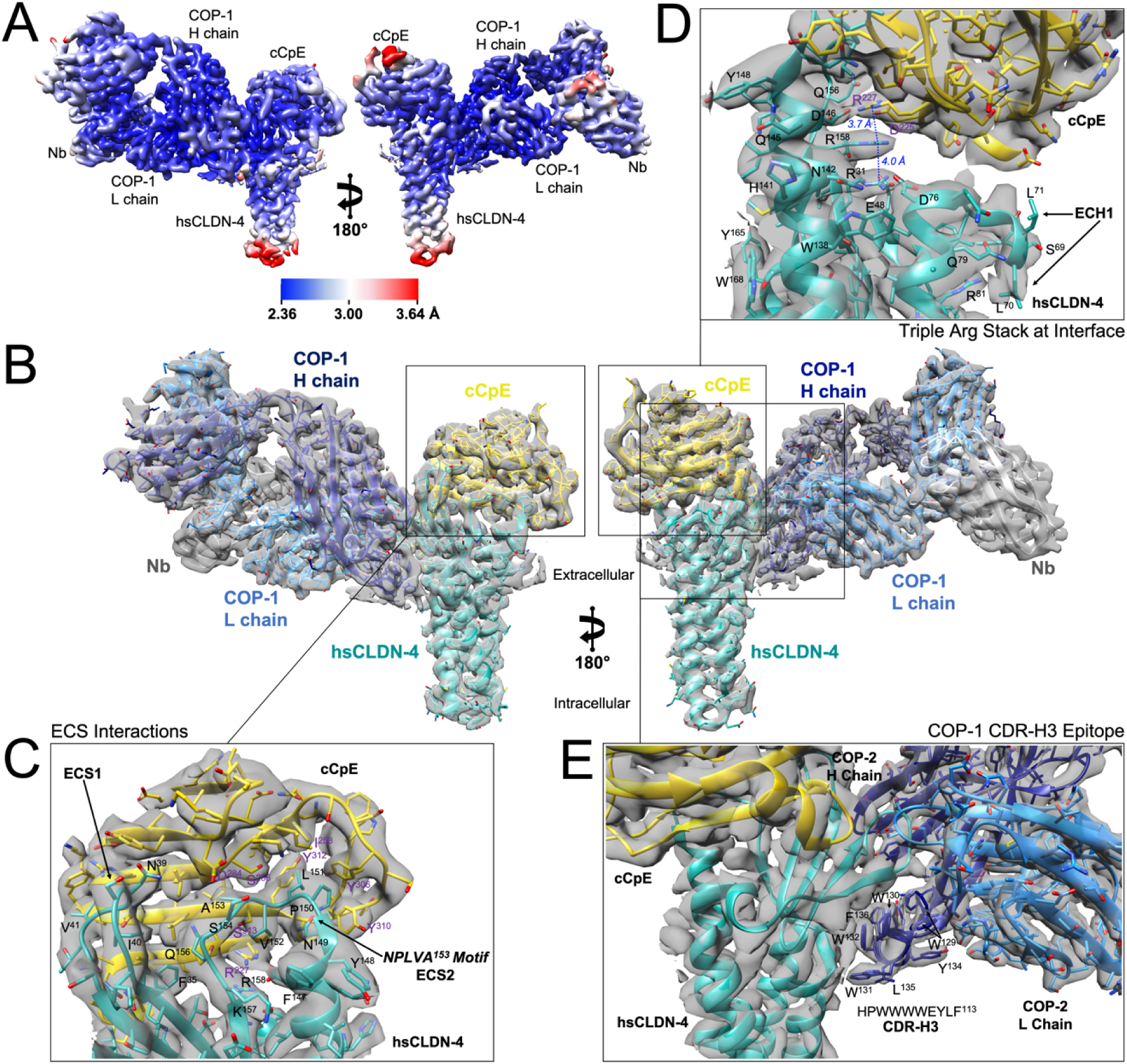
Structural Analysis of the Claudin-4/cCpE/COP-1/Nb Complex. (A) Local resolution estimates of the hsCLDN-4/cCpE/COP-1/Nb map. Regions of the map are colored according to highest (blue) and lowest (red) resolutions. (B) Global structure of the hsCLDN-4/cCpE/COP-1/Nb complex with overlaid map (grey). HsCLDN-4 (teal), cCpE (gold), COP-1 (blue) and NB (grey) are colored accordingly. (C) Zoom-in of the binding interface between the two ECS of hsCLDN-4 and cCpE. (D) Zoom-in of another binding interface between hsCLDN-4 and cCpE where three arginines form stacked interactions. (E) Zoom-in of the binding interface between hsCLDN-4 and the H chain (dark blue) of COP-1.

### Structural Similarity in hsCLDN-4/cCpE Complex Structures from Crystallography and Cryo-EM

We first analyzed the hsCLDN-4/cCpE portion of the complex because the resolution of this map exceeded electron density maps produced by crystallography from two existing structures of the hsCLDN-4/cCpE complex (3.4 and 3.5 Å). This was undertaken to decipher how COP-1 binding and/or the lack of crystal packing interactions affected interactions between hsCLDN-4 and cCpE reported previously.

Structural alignment of the hsCLDN-4/cCpE portion of the hsCLDN-4/cCpE/COP-1/Nb complex from cryo-EM and PDB ID 7KP4, the crystal structure of hsCLDN-4/cCpE, revealed a root mean square deviation of 0.90 Å between structures (***SI Appendix*, Fig. S3A**). Global structural elements and side chain orientations are largely unchanged, thus interactions between hsCLDN-4 and cCpE are preserved. Specifically, the NPLVA^153^ motif, which imparts high-affinity cCpE binding to claudins (**Fig. 3C**); as well as the triple arginine stack at the hsCLDN-4/cCpE interface that imparts claudin-selective binding to cCpE (**Fig. 3D**) [39]. The positions of these arginines are coordinated by acidic residues in a second shell. The Cζ-Cζ distances between hsCLDN-4 Arg158 and the other arginines are 3.7 and 4.0 Å in the cryo-EM structure, whereas in the crystal structure they were 3.5 and 3.8 Å. This structural overlay also showed that two extracellular loops (ECL) in ECS1 and the extracellular helix (ECH1) connecting /34–TM2 were conformationally different between structures (***SI Appendix*, Fig. S3B**). The conformations in the cryo-EM structure appeared to be altered due to COP-1 binding because COP-1 interacts with the two loops in ECS1 (***SI Appendix*, Fig. S3C**). CDR-L3 and CDR-H3 coordinate to sandwich ECL2 of hsCLDN-4 between them (**Fig. 3E** and ***SI Appendix*, Fig. S3C**). Although these COP-1-induced changes alter hsCLDN-4 ECS1 they do little to alter the global orientation of cCpE or perturb critical interactions between it and hsCLDN-4. In fact, COP-1 does not directly interact with cCpE (**Fig. 3B** and **3E**). Overall, the cryo-EM structure confirms the interactions and conformations hsCLDN-4 and cCpE utilize to engage in a high-affinity complex and validates the crystal structure, thus supporting conclusions made from that analysis.

### Deciphering the Extra-Membraneous Surface of hsCLDN-4

Unlike crystallography, detergent micelles are resolvable in cryo-EM maps (**Fig. 1D** and **1E**). We exploited this to visualize what claudin surfaces are accessible to sFabs and for inferring the boundary of the membrane. We first validated whether the cryo-EM density for the micelle correlated to a logical membrane boundary by inputting the structure into the Positioning Protein in Membranes (PPM) Server, a computational tool that energetically optimizes the spatial positions of membrane proteins in model membranes from structural coordinates [43]. PPM predicted a membrane that strongly correlated with the experimental density from the micelle, even though the micelle was not included as input (**Fig. 4A**). The inner and outer leaflet boundaries resided at the extremities of the micelle torus. The predicted membrane was validated further by the presence of strong density between COP-1s CDR-H3 and hsCLDN-4, which we modeled as a LMNG detergent due to the density’s “X” shape, which fit LMNGs branched chain structure (**Fig. 4A**). The boundary of the outer membrane leaflet ended at the point between the sugar headgroup and acyl chain of LMNG, which was also not input into the server (**Fig. 4B**). The membrane boundary estimated by PPM thus made physicochemical sense and could be visually validated by the density for LMNG. PPM predicted that 96 (46%) of hsCLDN-4s 209 amino acids were membrane-embedded, leaving 113 (54%) to reside outside the membrane and of these, 83 (40%) comprise a structured extracellular surface to target with sFabs. This analysis proved important because distinguishing the membrane boundary provided a marker to elucidate the basis of COP-1 binding to hsCLDN-4.

**Figure 4.**
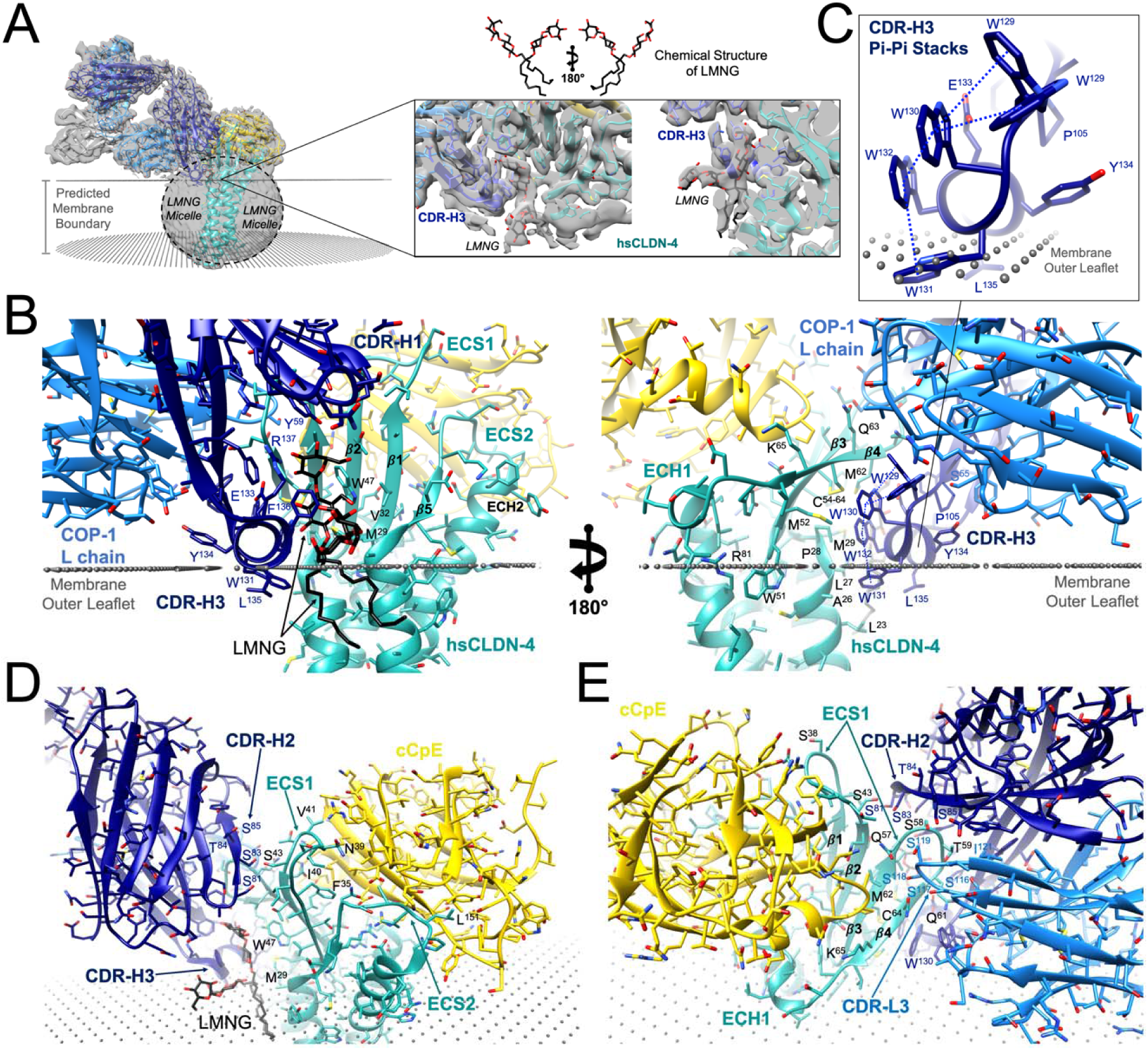
Structural Basis of COP-1 Recognition of Human Claudin-4. (A) Cryo-EM map from the hsCLDN-4/cCpE/COP-1/Nb complex (grey) contoured to show the experimentally determined boundary of the detergent micelle with predicted membrane boundary from the PPM server (black dots) [43]. Zoom-in of the interface between hsCLDN-4 (teal) and COP-1 (blue) showing a modelled LMNG detergent. The chemical structure of LMNG in a stick representation is also shown with carbons (black) and oxygens (red) colored accordingly. (B) The predicted PPM membrane boundary (black dots) overlaid on the structure of hsCLDN-4/cCpE/COP-1/Nb complex. HsCLDN-4 (teal), cCpE (gold), COP-1 (blue) and NB (grey) are colored accordingly, along with LMNG, which is colored as in A. (C) Zoom-in on the structure of COP-1s CDR-H3 amphipathic helix showing π-π stacking and associated membrane penetration. (D) Interactions between COP-1s H chain (dark blue) and hsCLDN-4 and LMNG. (E) Interactions between COP-1s H (dark blue) and L (light blue) chains with hsCLDN-4.

### Structural Basis of COP-1 Recognition of hsCLDN-4

COP-1s inability to significantly affect interactions between the hsCLDN-4/cCpE complex, we hypothesized, could be explained by its lack of interaction with cCpE. We therefore sought to decipher how COP-1 effectively targets and binds the complex using only hsCLDN-4, which had only a 40% by mass accessible antigenic surface. We analyzed COP-1s binding interface with hsCLDN-4 starting from the visualized micelle and estimated membrane boundary from PPM (**Fig. 4B**). The WWEYLF^136^ portion of COP-1s CDR-H3 forms an amphipathic llJ-helix that abuts and penetrates the detergent micelle, interacting with both hsCLDN-4 and LMNG (**Fig. 4A** and **4B**). Trp131 and Leu135 in the WWEYLF^136^ helix reside within the hydrophobic part of the micelle (**Fig. 4B**). Leu135 does not interact with hsCLDN-4 within the micelle but Trp131 does, forming non-polar interactions with Leu23, Ala26, and Leu27 on TM1 (**Fig. 4B**). Other residues of the WWEYLF^136^ helix make interactions with hsCLDN-4 outside of the membrane. Trp130, Trp132, and Glu133 form non-polar and polar interactions with hsCLDN-4; while Trp132, Glu133, and Phe136 also interact with LMNG (**Fig. 4B**). The helical structure of WWEYLF^136^ is facilitated by sequential π-π stacking interactions between Trp129 thru Trp132 in CDR-H3 where Trp129/130/132 form parallel-displaced and Trp131/132 T-shape stacks (**Fig. 4C**). The density for Trp129 showed that it has an alternate conformation that can parallel-displace or T-shape stack with Trp130. Trp129 dynamics may be due to Ser55 of CDR-L1, which resides 3.2 Å away and could hydrogen bond with it. Sequence alignments show that the PWWWWEYLF^136^ sequence of COP-1s CDR-H3 is not present in other COPs (***SI Appendix*, Fig. S1**). COP-1s ability to penetrate a hydrophobic environment gives it access to generally inaccessible TM regions, effectively increasing the accessible surface area on hsCLDN-4 for targeting.

Other regions and residues of COP-1 outside of CDR-H3 also form interactions of significance. Tyr59 and His61 of CDR-H1 interact with the /33–/34 loop of hsCLDN-4. In CDR-H2, Ser81, 83 and 85 interact with both ECLs of hsCLDN-4 (**Fig. 4D** and **4E**). In full, 10 residues of COP-1s H chain interact with two epitopes of hsCLDN-4 (***SI Appendix*, Table S1**). Epitope 1 constitutes residues LCCALPM^29^; epitope 2, QSTGQMQC^64^ of hsCLDN-4 (***SI Appendix*, Fig. S4A**). COP-1s L chain plays a more minor role than chain H with only six side chains making interactions with epitope 2 of hsCLDN-4 (**Fig. 4E** and ***SI Appendix*, Table S1**). Both COP-1 chains thus overlap interaction sites on hsCLDN-4 as they both bind epitope 2 (***SI Appendix*, Fig. S4B**). In chain L, CDR-L3 drives most interactions with Ser 116, 117, 118, and 119 forming up to 10 potential hydrogen bonds with hsCLDN-4 epitope 2 residues Gln57, Ser58, Thr59, Gln61, and Gln63 (**Fig. 4E** and ***SI Appendix*, Table S1**). In sum, COP-1 employs both chains and five of six CDRs to bind distinct portions of hsCLDN-4 that reside inside and outside of the membrane. We hypothesized that this property of COP-1 could confer it with high-affinity and isoform-selective binding to hsCLDN-4.

### Biophysical Characterization of COP-1

We next tested whether COP-1 could bind hsCLDN-4 selectively over other subtypes due to its distinct binding mode by determining the affinity and kinetics of its interactions. Using bio-layer interferometry (BLI), we established qualitative and quantitative binding analyses between claudins, cCpE, CpE, and COP-1 (***SI Appendix*, Fig. S5**). We first validated cCpE binding to human claudins -3 (hsCLDN-3), -4, and -9 (hsCLDN-9); and mouse claudins -3 (mmCLDN-3) and -4 (mmCLDN-4). These claudins were chosen due to their known receptor capacities for CpE and homologous sequences (***SI Appendix*, Fig. S6**). We determined rates for the second-order association (k_on_) and first-order dissociation (k_off_) constants, and equilibrium dissociation constant (K_D_), and found that all claudins except hsCLDN-3 bound with <12 nM affinities (**Table 1** and ***SI Appendix*, Fig. S5A-E**). These values agreed with our previously published values, although the affinity of cCpE to mmCLDN-4 was not reported [38, 39]. These results initiated in-depth analyses of COP-1 binding to claudin/enterotoxin complexes.

**Table 1.**
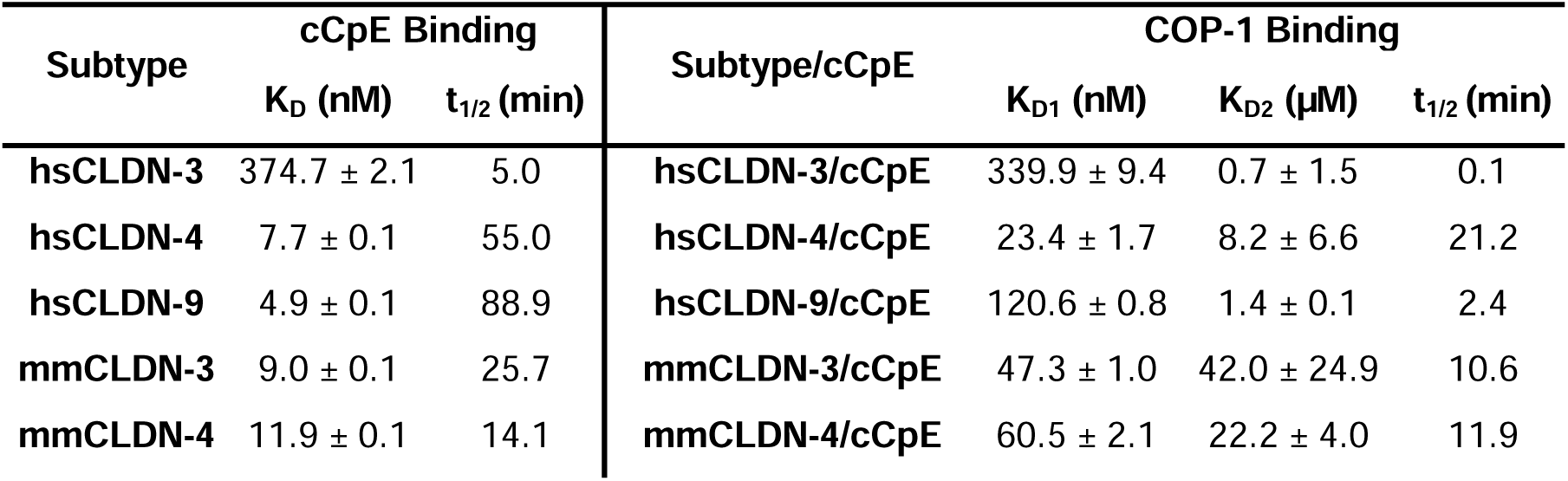
Binding of Claudins to cCpE and COP-1 to Claudin/cCpE Complexes. BLI was used to measure binding kinetics and affinities. Binding of claudins to cCpE represent a single experiment and was fit with a 1:1 binding model, while COP-1 binding measurements represent averages from duplicate experiments and were fit with a 2:1 heterogenous ligand model, hence two K_Ds_ are reported. The t_1/2_ (ln 2/k_off_) was calculated from the dissociation rates between claudin/cCpE and COP-1 that represent K_D1_. The full binding results, including kinetic rates, appear in ***SI Appendix*, Table S2**, with experimental details provided in **Materials and Methods, Biophysical Characterization**.

Once it was established that all claudins bound to cCpE we qualitatively assessed claudin binding to CpE; COP-1 binding to cCpE; and COP-1 binding to claudins using single-point analyses (***SI Appendix*, Fig. S5F-H**). We found that all claudins bound CpE with a hierarchy of hsCLDN-4>hsCLDN-9>mmCLDN-3>mmCLDN-4>>hsCLDN-3 (***SI Appendix*, Fig. S5F**). These results mirrored those from binding to cCpE, with hsCLDN-3 binding more weakly than other subtypes. We then determined that COP-1 did not bind to cCpE (***SI Appendix*, Fig. S5G**). This result verified our biochemical assay where we found that excess COP-1 and cCpE did not elute as a complex (**Fig. 1A**). We next found that COP-1 did bind to claudins in absence of cCpE, and had a strong binding preference for hsCLDN-4, followed by mmCLDN-4>hsCLDN-3>mmCLDN-3>hsCLDN-9 (***SI Appendix*, Fig. S5H**). These analyses made it clear that COP-1 binds to claudins but not cCpE, so we then measured COP-1 binding to claudin/enterotoxin complexes as this knowledge was relevant to understand the structures.

### Biophysical Basis of COP-1 Selectivity

To qualitatively grasp COP-1 binding to claudin/enterotoxin complexes we performed single-point analyses (***SI Appendix*, Fig. S5I-J**). Here, pre-formed claudin/cCpE were immobilized then binding was tested against COP-1. Results showed that COP-1 bound complexes in a subtype-specific way with a preference for hsCLDN-4/cCpE>mmCLDN-3/cCpE>mmCLDN-4/cCpE>>hsCLDN-9/cCpE—COP-1 did not bind hsCLDN-3/cCpE (***SI Appendix*, Fig. S5I**). We then tested COP-1 binding to claudin/CpE, where CpE contains an additional cytotoxic N-terminal domain in addition to its cCpE domain. We found that COP-1 bound with identical preferences to claudin/CpE as to claudin/cCpE—again COP-1 failed to bind hsCLDN-3/CpE (***SI Appendix*, Fig. S5J**). Because k_on_ and k_off_ rates were measured, the single-point binding curves could be fit to a binding model to generate an approximate K_D_. Interestingly and unlike claudin binding to enterotoxins, COP-1 binding data best fit to a 2:1 heterogeneous ligand model. In sum, these results showed that COP-1 was selective for hsCLDN-4/enterotoxin complexes despite sequence identities between subtypes that ranged from 65.7% hsCLDN-9, 68.3% mmCLDN-3, 69.2% hsCLDN-3, to 83.7% mmCLDN-4 (***SI Appendix*, Fig. S6**).

Although the qualitative binding studies provided insights into the binding mode and selectivity of COP-1, we conducted complete quantitative studies with claudin/cCpE to determine the kinetics and affinities of COP-1 binding. Here, using pre-formed claudin/cCpE complexes, we tested binding against six concentrations of COP-1 to hsCLDN-3/cCpE (***SI Appendix*, Fig. S5K**), hsCLDN-4/cCpE (***SI Appendix*, Fig. S5L** and **S5P**), hsCLDN-9/cCpE (***SI Appendix*, Fig. S5M**), mmCLDN-3/cCpE (***SI Appendix*, Fig. S5N**), and mmCLDN-4/cCpE (***SI Appendix*, Fig. S5O**). This data also fit best to a 2:1 heterogeneous ligand binding model and thus two K_Ds_ are reported, one for each of the estimated two COP-1 interactions. We found that COP-1 bound with an average K_D1_ of 339.9, 23.4, 120.6, 47.3, and 60.5 nM; and an average K_D2_ of 0.7, 8.2, 1.4, 42.0, and 22.2 µM, respectively (**Table 1**). The hierarchy of COP-1 K_D1_ affinity was hsCLDN-4/cCpE>mmCLDN-3/cCpE>mmCLDN-4/cCpE>hsCLDN-9/cCpE>hsCLDN-3/cCpE, which was in accordance with the single-point results (***SI Appendix*, Fig. S5I**). In this experiment, COP-1 bound to hsCLDN-3/cCpE because concentrations above its K_D1_ were used. Additionally, we used these more accurate k_off_ rates to calculate the half-life (t_1/2_) of claudin/cCpE/COP-1 complexes, as this time measurement provides added context to affinity measurements. The hierarchy of COP-1 half-lives were hsCLDN-4/cCpE>mmCLDN-4/cCpE>mmCLDN-3/cCpE>hsCLDN-9/cCpE>hsCLDN-3/cCpE (**Table 1**). These findings provided more accurate measurements by which to elucidate the biophysical and mechanistic bases of COP-1s targeting and selective binding to claudins with high homologies.

### Biochemical Validation of COP-1 Selectivity

Lastly, we sought to determine whether COP-1-bound claudin/cCpE complexes could be retained through SEC, which if so, would validate the biophysical findings. Here, pre-formed claudin/cCpE complexes were removed from wells of the 96-well plate used for BLI and injected on an SEC column. Another aliquot of these complexes was incubated with COP-1 and injected on SEC. This experiment showed that in the absence of COP-1, all claudin/cCpE samples elute between 10.25-10.60 minutes (***SI Appendix*, Fig. S7A**). When COP-1 is added, larger molecular weight complexes were formed by hsCLDN-4/cCpE, mmCLDN-3/cCpE, and mmCLDN-4/cCpE, as determined by decreased peak elution times <10 minutes (***SI Appendix*, Fig. S7B**). For hsCLDN-9/cCpE, two peaks eluted, one <10 minutes and another at 10.45 minutes. For hsCLDN-3/cCpE, the peak elution time with COP-1 was 10.40 minutes, similar to 10.60 minutes without COP-1. To validate that the peaks eluting at <10 minutes were COP-1-bound we added COP-2—a cCpE-binding sFab that binds claudin/cCpE complexes—to hsCLDN-4/cCpE and found this complex eluted at 9.50 minutes (***SI Appendix*, Fig. S7B**). This experiment showed that COP-1 bound to hsCLDN-4, mmCLDN-3, and mmCLDN-4, while hsCLDN-9 formed a partial complex and hsCLDN-3 did not bind COP-1 (***SI Appendix*, Fig. S7C**). This experiment validated our biophysical results by showing which claudin/cCpE complexes bind COP-1 with sufficient affinity to remain intact throughout SEC. Coupled with the structural results, our biochemical and biophysical findings reveal COP-1s basis for selectivity to hsCLDN-4 and provides a framework to use sFabs to bolster structural and therapeutic efforts that target claudins.

## Discussion

We have developed a platform using hsCLDN-4/cCpE complexes and a phage display library encoding sFabs that yields selective binding molecules and molecular structures of claudins. We previously showed that the cCpE binding sFabs COP-2 and COP-3 can enable low resolution structures of claudins by cryo-EM at 200 kV. Now, using the claudin-binding sFab COP-1 and 300 kV microscopes, we demonstrate that COPs can enable high resolution cryo-EM structures of 22 kDa claudin-4 and its complex with a 15 kDa enterotoxin fragment from *Clostridium perfringens*. The 2.6 Å resolution achieved using this platform surpasses X-ray crystallographic structures of this complex and the resulting cryo-EM structure validates the crystallographic structure by showing that the intermolecular interactions observed between hsCLDN-4 and cCpE are conserved in both. The consistency in structures determined by two independent methods therefore provides verification that the NPLVA^153^ and cCpE-binding motifs of claudins direct recognition and high-affinity binding by CpE, which we exploited to develop the sFab-driven platform [39]. The discovery of a claudin-binding sFab and the structures conferred by it provide new strategies for targeting claudins and for modifying cCpE and CpE to modulate its claudin-selective binding properties. These insights can be applied to advance the development of therapeutics to tune permeability of the blood-brain barrier or visualize and destroy claudin over-expressing cancers, two current applications.

The cryo-EM structures reveal the basis of COP-1 binding to claudins. They show that COP-1s CDR-H3 uses a string of four sequential Trp residues that π-π stack to form an amphipathic helix that abuts and penetrates detergent micelles, giving COP-1 access to bind the hydrophobic TM regions of claudins. Other CDRs on the L and H chain contribute to COP-1 recognition and binding to claudin-4, which do so without affecting claudin-4s native interactions with cCpE. Because the interactions between hsCLDN-4 and cCpE are unaltered upon COP-1 binding, COP-1 appears to conformationally react to the pre-existing hsCLDN-4/cCpE complex rather than proactively changing it to bind its epitopes. In this way, COP-1 is poised to selectively bind the conformation that claudins are in when bound to cCpE. This is validated by our results showing that the hsCLDN-4/cCpE portion of the cryo-EM structure overlays very well onto the crystal structure (***SI Appendix*, Fig. S3**), and that COP-1 binds better to claudin/cCpE complexes versus claudins alone (***SI Appendix*, Fig. S5**).

Further evidence that COP-1 is conformationally reactive to claudin/cCpE complexes is provided by COP-1 binding curves, which fit best to a 2:1 model for heterogeneous ligands. We surmise that this is true because either COP-1 requires time to conformationally alter the structure of its CDR-H3 to sense, penetrate the membrane, and bind claudins; or BLI can distinguish the individual binding events of COP-1s H and L chains to claudins. This 2:1 binding mode exists for COP-1 binding to claudins alone or to claudin/cCpE complexes, which is further evidence that cCpE binding does not largely affect the binding mechanism of COP-1 to claudins. In addition to CDR-H3 of COP-1, the second ECL of hsCLDN-4 ECS1 binds between the H and L chains of COP-1. Thus, other movements could also account for this binding behavior. Further characterization can uncouple which explanation is experimentally valid. Either way, this finding shows that the process of COP-1 binding is complex and may rely on conformational changes within COP-1 but not its targeted claudin/cCpE epitope, which is why the mathematical fitting models demonstrate this heterogenous behavior. Other findings reported here provide evidence that COP-1 is isoform-selective as well as conformationally reactive.

Our biophysical results show that COP-1 binds claudins in an isoform-selective way and reveals the mechanism of its selectivity. In the absence of cCpE, COP-1 binds preferentially to hsCLDN-4, with mmCLDN-4, hsCLDN-3, mmCLDN-3, and hsCLDN-9 binding sequentially worse. This verifies that COP-1 is selective toward claudin-4 orthologs, which are 83.7% identical, over the other three subtypes, which exhibit <70% sequence identities compared to hsCLDN-4 (***SI Appendix*, Fig. S6**). At 65.7%, hsCLDN-9 has the lowest sequence conservation, which explains its decreased binding to COP-1 of all subtypes tested. The data from BLI also shows that the k_off_ rates of COP-1 binding to claudins is faster than when cCpE is bound (***SI Appendix*, Figure S5H-I**). This shows that COP-1 epitopes on claudins is likely dynamic in absence of cCpE. Binding of cCpE to claudins limits these dynamics and in turn, stabilizes COP-1 binding epitopes, resulting in longer complex association times compared to claudin alone. In this way, cCpE boosts COP-1s efficacy for binding claudins in an indirect way as no direct interactions between cCpE and COP-1 are observed. These results show that COP-1 may hold promise as a claudin binding and tight junction modulating molecule as it does not rely completely on cCpE to recognize and bind hCLDN-4.

When claudins are bound to cCpE, COP-1 binding selectivity appears to be altered slightly. Here, COP-1 binds preferentially to hsCLDN-4 as before, binds mmCLDN-3 and mmCLDN-4 well and hsCLDN-9 poorly, but binds hsCLDN-3 very poorly. Why, when bound to cCpE, is hsCLDN-3 now an inferior target of COP-1? The answer reflects a limitation of the assay where we use cCpE to immobilize the claudin/cCpE complex to the BLI sensor. Of all claudins tested, hsCLDN-3 has the lowest affinity for cCpE because of its rapid k_off_ and short t_1/2_ (**Table 1** and ***SI Appendix*, Table S2**). This means that the complex is not retained on biosensors long enough to detect COP-1 binding to hsCLDN-3 because only cCpE is bound to sensors. When taking this explanation for COP-1s low affinity for hsCLDN-3/cCpE into consideration, it appears clear then that COP-1 binds claudins in practically the same isoform-selective order whether they are unbound or bound to cCpE. Because COP-1 binds in a subtype-specific manner to claudins in the absence or presence of cCpE and CpE (***SI Appendix*, Table S5H-J**), it has added value as a molecular tool to probe or modify claudin structure/function under normal or CpE-induced pathogenic conditions.

Is COP-1 selectivity governed by recognizing unique conformations of claudins or by sequence divergence in key COP-1 binding epitopes of claudins? Our findings suggest the latter. Whether unbound or bound to cCpE, hsCLDN-9 exhibits the worst binding to COP-1 despite having high affinity and slow k_off_ rates for cCpE—yet it also has the lowest sequence identity to hsCLDN-4 of the subtypes assayed. This suggests that COP-1s poor binding to it reflects a lower selectivity due to sequence changes. But, we can use existing crystal structures to discern whether conformational differences between claudin/cCpE complexes drives COP-1 selectivity, as structures of hsCLDN-4, hsCLDN-9, and mmCLDN-3 bound to cCpE have been determined [37–39]. Comparison of the cCpE-bound hsCLDN-4, hsCLDN-9, and mmCLDN-3 structures show differences, but these are generally subtle alterations to ECS loops while major structural elements remain unchanged (***SI Appendix*, Fig. S8**). One significant difference is the conformation of cCpE when bound to mmCLDN-3, which is separated from the standard claudin/cCpE interface and resides in a more open conformation (***SI Appendix*, Fig. S8A**). Despite mmCLDN-3 having more conformationally flexibility, our biophysical evidence shows that it is hsCLDN-9 and not mmCLDN-3 that COP-1 binds poorly to. Because the conformations of hsCLDN-9/cCpE are like hsCLDN-4/cCpE, it suggests that sequence divergence between these two subtypes is what drives COP-1s selective binding to hsCLDN-4. This is supported by mmCLDN-3/cCpE, which is conformationally altered but more sequentially homologous and binds COP-1 well. This analysis suggests that the conformational changes observed in the mmCLDN-3/cCpE structure either do not affect COP-1 binding or are an artifact of crystal packing interactions and do not occur in solution. In light of the structural and biophysical results presented here with COP-1, this data supports a theory where cCpE binding does not largely alter the structure of claudins—any conformational changes that do occur are likely localized and/or minute and thus well-tolerated by COP-1. In this way, sFabs like COP-1 could be powerful probes of claudin structure as they could be engineered to sense slight conformational changes that occur upon cCpE binding or claudin self-assembly. Taken further, we postulate that our data provides experimental evidence that explains cCpEs mechanism for dissociating tight junctions [20]. If claudin structure is minimally altered by cCpE binding it suggests that cCpE-induced tight junction dissociation results from disrupting the equilibrium of claudin integration into tight junctions rather that physically breaking integrated claudin oligomers within intact tight junctions. Our findings support the “sequestering” model of cCpE disassembly of tight junctions proposed by Sonoda *et al.* [20].

If COP-1 selectivity is driven by sequence divergence in claudins, which residues direct COP-1 recognition? We show that the sequence identity between hsCLDN-4 to other subtypes used in this study range from 65.7% for hCLDN-9 to 83.7% for mCLDN-4 (***SI Appendix*, Fig. S6**). In the two epitopes COP-1 interacts with on hsCLDN-4 (***SI Appendix*, Table S1**), the sequence identity between claudins is 100% in epitope 2, and 85.7% between hsCLDN-4 and all of hsCLDN-3, mmCLDN-3 and mmCLDN-4—whereas it is 57.1% between hsCLDN-4 and hsCLDN-9. Since all subtypes have identical epitope 2s, this suggest epitope 1 is the driver of COP-1 recognition and binding. Epitope 1 contains seven consecutive amino acids that constitute the top of TM1 (***SI Appendix*, Fig. S4**). The first five residues reside in the membrane and last two are extracellular (**Fig. 4**). The most variable side chains in epitope 1 between subtypes are Leu23, Cys24, and Met29 (***SI Appendix*, Fig. S6**). We hypothesize that the substitutions of Val23 and Leu29 in hsCLDN-9 cause low COP-1 affinity because they face CDR-H3 and, due to the shortened lengths of these hydrophobic side chains compared to Leu23/Met29 from hsCLDN-4, disrupt interactions with Trp131 and Trp132 in COP-1s amphipathic CDR-H3. Because no other subtype tested shares 100% sequence identity with hsCLDN-4 in epitope 1 they all bind COP-1 worse due to substitutions of their corresponding Leu23 or Met29 (**Table 1**). We anticipate that these side chains and epitope 1 in general are major drivers for COP-1-selective binding of claudins. Sequence alignments of 23 human claudins reveals that the LCCALPM^29^ motif is highly specific to hCLDN-4 alone, with no other subtype having this sequence [38]. In fact, only hsCLDN-3, -5, -6, and -9 share identities in epitope 1 above 57%—and we have shown that COP-1 can distinguish hsCLDN-4 from -3 and -9. Based on sequence analysis, we predict that COP-1 binds human claudins with the following selectivity (from best to worse): hsCLDN-4, -3, -5 and -6, and finally, -9. This idea requires experimental validation, but if validated, provides proof-of-principle that COP-1 or similarly functioning sFabs can selectively target claudins, be used to alter their cis or trans interactions, and in turn, alter the permeabilities of tight junctions in potentially subtype- and/or tissue-specific ways.

## Summary

We have developed, characterized the binding, and determined a structure of a sFab called COP-1 and confirmed its selective binding for human claudin-4 over other homologous human and mouse subtypes. The structure and corresponding analyses provide the structural and mechanistic bases of COP-1s subtype-selective recognition and binding of human claudin-4 and proof-of-principle that it or similarly engineered molecules could be used to target specific claudins, which in turn may be used to tune the molecular permeability of tight junctions in controlled ways. To date, few molecules in the literature have been developed and experimentally validated to bind directly to claudins, and none have had their structures determined in complex with a claudin. This work thus enhances insights and strategies for effective molecular targeting of claudins inside and outside the membrane. Moreover, it highlights the advantages of using sFabs for these purposes as they are easy to produce, monovalent, stable, inherently bind with high affinity and selectivity, and can be rationally optimized to improve these properties. These traits provide an added benefit as we also demonstrate that sFabs enable cryo-EM structure determination of a 22 kDa membrane protein, eliminates the bottleneck of crystallization, yet still resolves structures of these small proteins with high precision. In full, the platform developed here has potential to serve both experimentalists and clinicians alike that are interesting in elucidating the form and function of tight junctions driven by claudins.

## Materials and Methods

### Expression, Purification, and Labeling of Claudins and Enterotoxins

All claudins (hsCLDN-3, -4, -9, mmCLDN-3 and -4) and enterotoxins (cCpE and CpE) were expressed and purified as described previously [39, 40]. Briefly, all proteins contained a C-terminal decahistidine tag preceded by a thrombin cleavage site (claudin-_His10_, cCpE-_His10_, and CpE-_His10_) and were expressed in insect cells. After cell lysis, membranes were prepared for claudins by ultracentrifugation while the supernatant was used for purification of enterotoxins. Claudins were solubilized from membranes using n-dodecyl-β-D-maltopyranoside (DDM, Anatrace) and cholesteryl hemisuccinate Tris salt (CHS, Anatrace) to a final concentration of 1/0.1% (w/v). After ultracentrifugation at 100,000 xg, the supernatants were used for purification by adding NiNTA resin (ThermoFisher) to them. For most applications, the protein-bound resin was washed and proteins were released and collected from the resin by digestion with thrombin. For enterotoxins, no detergents were used while for claudins, the concentrations of DDM/CHS in the washes were 0.087/0.0087%, and 0.04/0.004% for protein release. All proteins isolated using these methods were pure as assessed by SDS-PAGE and analytical size-exclusion chromatography (SEC) using a Superdex 200 increase column equilibrated in SEC Buffer (20 mM Hepes pH 7.4, 100 mM NaCl, 1% glycerol). For claudins, SEC buffer also contained 0.04% DDM, with no CHS. These tagless post-NiNTA proteins were frozen in liquid nitrogen at 1 mg/mL and stored at -80°C until use.

For structural studies, hCLDN-4 was solubilized using 1/0.1% DDM/CHS, ultracentrifuged, and bound to NiNTA resin as above. However, during NiNTA purification, it was exchanged from DDM/CHS to 2,2-didecylpropane-1,3-bis-β-D-maltopyranoside (LMNG, Anatrace)/CHS by collecting the resin with a bead-capture column (Bio-Rad) then washing it for 10 column volumes in Wash Buffer A (50 mM Tris pH 7.4, 500 mM NaCl, 20 mM imidazole, 10% glycerol, and 0.1/0.01% LMNG/CHS). Captured resin was washed for another 10 column volumes in Wash Buffer B (50 mM Tris pH 7.4, 300 mM NaCl, 40 mM imidazole, 5% glycerol, and 0.1/0.01% LMNG/CHS). Beads were captured and washed for two column volumes with Cleavage Buffer (50 mM Tris pH 8.0, 150 mM NaCl, and 0.05/0.005% LMNG/CHS). After washing, 3 mL of Cleavage Buffer were added to resin and hCLDN-4 was released by thrombin overnight. The next day, hCLDN-4 in LMNG/CHS were captured in the flow-through, analyzed by SDS-PAGE and analytical SEC, and used to form complexes, which is detailed further in **Preparation of Complexes for Cryo-EM**.

For biophysical studies, claudins and/or enterotoxins required immobilization on biosensors. For claudins, the tagless post-NiNTA pure samples were biotinylated while enterotoxins were eluted off NiNTA using imidazole to maintain their C-terminal decahistidine tags. Briefly, claudins were purified as above and equilibrated in Biotin buffer (20 mM Hepes pH 7.2, 150 mM NaCl, 1% glycerol, and 0.04% DDM). This pH specifically labels the N-terminal amine of claudins using NHS chemistry. Claudins were labeled with NHS-PEG4-Biotin (ThermoFisher) using 4:1 biotin:claudin molar excess overnight at 4°C. The next day, 50 mM Tris pH 7.4 was added to quench the reaction and the samples were buffer exchanged into BLI buffer (20mM Tris pH 7.4, 100mM NaCl, 1% glycerol, and 0.03% DDM) using PD-10 columns (Bio-Rad). For enterotoxins, the proteins were released from NiNTA resin after incubation with Elution buffer (50 mM Tris pH 7.4, 150 mM NaCl, 300 mM imidazole, and 1% glycerol). Enterotoxins were then buffer exchanged into BLI buffer using a PD-10 column to remove imidazole. All claudin-_Biotin_ and cCpE-_His10_ and CpE-_His10_ proteins were analyzed by SDS-PAGE and analytical SEC and judged to be biochemically homogenous and pure; at which point they were frozen in liquid nitrogen at 1 mg/mL and stored at -80°C until use.

### Development, Validation, Expression and Purification of COP-1

COP-1 was developed using a phage display library encoding synthetic antigen-binding fragments (sFabs) concurrent with previously reported sFabs COP-2 and COP-3 [40]. Preparation of hsCLDN-4/cCpE complexes are provided therein. For panning, DDM-solubilized and biotinylated hCLDN-4 in complex with cCpE was used as input into the phage display pipeline after polishing and removing free biotin using SEC. COP-1 was therefore developed, validated, expressed, and purified identically to previously described methods [40]. During purification, COP-1 was found to be less soluble in buffers than COP-2 and -3, and so 0.01% DDM was added during affinity purification. After affinity purification, COP-1 was exchanged to BLI buffer, frozen in liquid nitrogen, and stored at -80°C until use in biochemical, biophysical, and structural biology studies.

### Preparation of Complexes for Cryo-EM

HsCLDN-4/cCpE complexes were formed using hCLDN-4 in LMNG/CHS (250 µg) and 1.2 molar excess of tagless cCpE incubated at 4°C for 1 hour. COP-1 was then added at a 1.0 molar ratio to hCLDN-4 and incubated with the hCLDN-4/cCpE complex at 4°C overnight. The next day, 1.5 molar excess of V_H_H nanobody (Nb) to COP-1 was added and incubated at 4°C overnight. The assembled hCLDN-4/cCpE/COP-1/Nb complexes were then concentrated using a 100 kDa MWCO concentrator (Millipore), 0.2 µm filtered, then loaded onto a Superdex 200 increase 10/300 column equilibrated in Cryo buffer (20 mM Hepes pH 7.4, 100 mM NaCl, and 0.003% LMNG). Fractions from SEC were run on SDS-PAGE and those containing the hCLDN-4/cCpE/COP-1/Nb complex were pooled and concentrated to 7 mg/mL using a 100 kDa MWCO concentrator. A fraction of this sample was also diluted in half to 3.5 mg/mL using Cryo buffer so that two concentrations could be tested in vitrification. Samples were shipped to the University of Chicago overnight on ice for vitrification and cryo-EM analyses.

Samples of a hCLDN-4/cCpE/COP-1 complex lacking the Nb were also prepared as described above. This complex was pooled and concentrated to 6 mg/mL then shipped overnight on ice to the Pacific Northwest Cryo-EM Center (PNCC) for vitrification and cryo-EM analyses.

### Cryo-EM Vitrification, Data Collection and Processing

Once received, samples of the hCLDN-4/cCpE/COP-1/Nb complex were placed on ice and vitrified the same day. Grids for cryo-EM analyses were glow-discharged for 60 seconds at 15 mA in a Pelco easiGlow (Ted Pella Inc) instrument then vitrified using a Vitrobot Mark IV (ThermoFisher) plunge freezing apparatus. Aliquots (3 μL) of hCLDN-4/cCpE/COP-1/Nb at both 7 and 3.5 mg/mL were each applied to a Quantifoil R1.2/1.3 200 mesh grid, an Au-Flat 0.6/1.0 300 mesh grid, and a UltrAuFoil 0.6/1.0 300 mesh grid at 4°C and 100% relative humidity. Grids were blotted for 5 seconds with a blot force of 2 and plunge frozen into liquid ethane cooled by liquid nitrogen. Grids were stored in liquid nitrogen before imaging. The six grids were screened for thin ice and particle presence and distribution. The 7 mg/mL sample frozen on UltrAuFoil 0.6/1.0 300 mesh grid was used for data collection.

Cryo-EM data collection for the hCLDN-4/cCpE/COP-1/Nb complex was performed on a Titan Krios G3i (ThermoFisher) equipped with a Gatan K3 direct electron detector and BioQuantum GIF at the University of Chicago Advanced Electron Microscopy Core Facility (RRID:SCR_019198). 5,039 movies were collected using EPU (ThermoFisher) in CDS mode at 81,000× magnification with a super resolution pixel size of 0.534 Å and physical pixel size of 1.068 Å, defocus range of 0.9 to 2.1 μm using a step of 0.2 μm, with a total dose of 60 electron/Å^2^ fractionated over 50 total frames.

Once the hCLDN-4/cCpE/COP-1 complex was received, it was also placed on ice then vitrified the same day. Grids for cryo-EM analyses were glow-discharged for 60 seconds at 15 mA in a Pelco easiGlow (Ted Pella Inc) instrument then vitrified using a Vitrobot Mark IV (ThermoFisher) plunge freezing apparatus. Aliquots (3 μL) of hCLDN-4/cCpE/COP-1 were applied to a Quantifoil R2/1 200 mesh grid at 4°C and 100% relative humidity. Grids were blotted for 1.5 seconds with a blot force of 2 and plunge frozen into liquid ethane cooled by liquid nitrogen. The grid were stored in liquid nitrogen prior to screening and data collection.

Cryo-EM data collection for the hCLDN-4/cCpE/COP-1 complex was performed on a Titan Krios G3i (ThermoFisher) equipped with a Gatan K3 direct electron detector and BioContinuum GIF at PNCC. 1,073 movies were collected using SerialEM in counting mode at 92,000× magnification with a super resolution pixel size of 0.2535 Å, defocus range of 0.8 to 2.2 μm using a step of 0.2 μm, with and a total dose of 60 electron/Å^2^ fractionated over 50 total frames.

All micrograph and particle processing for both complexes were performed in CryoSPARC [44] . Patch-motion correction and patch-CTF correction were used to correct for beam-induced motion and calculate CTF parameters from the motion-corrected micrographs. Blob-based template picking followed by 2D classification were used to generate templates that were subsequently used for template-based particle picking. Particles identified from this template-based picking procedure were subjected to at least one round of 2D classification, followed by *ab initio* 3D reconstruction, heterogeneous and homogenous refinement, and finally non-uniform and local refinement. A workflow is show in ***SI Appendix*, Fig. S9**.

### Model Building, Refinement and Structure Determination

A 3D model of COP-1 was generated using AlphaFold2 (AlphaFold2_advanced.ipynb) [45]. Then, the crystal structure of hCLDN-4 bound to cCpE (PDB ID 7KP4) and the COP-1 model were independently placed into the cryo-EM density manually then fit into the map with Chimera [46]. The sFab/Nb complex from PDB ID 7ZLG that contains a sFab L chain-binding V_H_H nanobody [42] was then superposed onto COP-1 by aligning the L chains of the sFab and COP-1. The 3D coordinates of the Nb and COP-1 were then written into a .pdb file containing this orientation, as well as the fit model of hCLDN-4/cCpE from the crystal structure, creating a file with all four proteins and their respective five chains. Each chain was then rigid body and real space refined into the cryo-EM map using Coot [47]. Refinement of the model was done iteratively using Namdinator and phenix.real_space_refine to optimize model-to-map fit, resulting in the final structure of the hCLDN-4/cCpE/COP-1/Nb complex [48, 49]. The hCLDN-4/cCpE/COP-1 complex structure was built using the claudin-4/cCpE/COP-1/Nb complex after deleting the Nb from the model. The programs used to visualize and build the structures included Coot and Chimera, refinement in Phenix, and figures were made using Chimera—using the SBGrid Consortium Software Suite [46–48, 50]. ***SI Appendix*, Table S4** shows data collection, refinement, and validation statistics for the structures.

### Biophysical Characterization

For biophysical analyses, we used claudin-_Biotin_, tagless cCpE, cCpE-_His10_, and CpE-_His10_ and post-affinity-purified COP-1 all solubilized in BLI buffer (20mM Tris pH 7.4, 100mM NaCl, 1% glycerol, and 0.03% DDM). Bio-layer interferometry (BLI) was performed at 25°C in 96-well black flat bottom plates (Greiner) using an acquisition rate of 5 Hz averaged by 20 using an Octet© R8 BLI System (FortéBio/Sartorius), with assays designed and setup using Blitz Pro 1.3 Software. Binding experiments consisted of the following steps: sensor equilibration (30 seconds), loading (120 seconds), baseline (60 seconds), and association and dissociation (200-300 seconds each).

The first experiments consisted of validating claudin binding to cCpE. Here, 500 nM of cCpE-_His10_ was immobilized on NiNTA (Dip and Read) sensors and associated with 0-600 nM claudin-_Biotin_ (***SI Appendix*, Fig. S5A-E**). Association and dissociation were done for 300 seconds to compare to previously reported results [38, 39]. We next tested CpE binding to claudins where 500 nM of CpE-_His10_ was immobilized on NiNTA sensors and associated with 200 nM claudin-_Biotin_ (***SI Appendix*, Fig. S5F**). These and all subsequent associations and dissociations were done for 200 seconds. For cCpE binding to COP-1, 500 nM of cCpE-_His10_ was immobilized in NiNTA biosensors and binding was tested against 0-750 nM COP-1 (***SI Appendix*, Fig. S5G**). For COP-1 binding to claudins, 1 µM of claudin-_Biotin_ was immobilized on Streptavidin-SA (Dip and Read) biosensors and binding was tested against 500 nM COP-1 (***SI Appendix*, Fig. S5H**). General trends for COP-1 binding to claudin/enterotoxin complexes were then performed as follows. First, 300 nM cCpE-_His10_ was incubated with 300 nM of claudin-_Biotin_ at 20°C for 1 hour to make complexes and then this 600 nM of pre-formed claudin-_Biotin_/cCpE-_His10_ complexes were immobilized on NiNTA biosensors and binding was tested against 250nM COP-1 (***SI Appendix*, Fig. S5I**). This same method was used to test COP-1 (250 nM) binding to 600 nM pre-formed claudin-_Biotin_/CpE-_His10_ complexes (***SI Appendix*, Fig. S5J**). Once general trends were established, we obtained the full binding and kinetic constants of claudin/cCpE against COP-1 as follows: 600 nM pre-formed claudin-_Biotin_/cCpE-_His10_ complexes were immobilized on NiNTA sensors then associated with 0-750 nM COP-1 (***SI Appendix*, Fig. S5K-O**). This strategy was chosen over immobilizing pre-formed claudin-_Biotin_/cCpE-_His10_ complexes on SA sensors after it was determined that NiNTA sensors resulted in greater signal strength. Here, 1,200 nM of pre-formed hCLDN-4-_Biotin_/cCpE-_His10_ complexes were immobilized on SA biosensors and binding was tested against 0-750 nM COP-1, which resulted in similar dissociation constants compared to those obtained from NiNTA sensors (***SI Appendix*, Fig. S5P**). Immobilizing half the concentration of hCLDN-4/cCpE complex resulted in ∼5-fold higher binding signal. We attribute this to the relatively small NiNTA moiety compared to the larger streptavidin (52 kDa), which results in more complex bound per sensor and thus greater light interference upon COP-1 binding. Mutant claudin-_Biotin_ experiments were performed identically using the NiNTA strategy with pre-formed cCpE-_His10_ complexes. Results for multi-point quantitative analyses appear in ***SI Appendix*, Fig. S5** and **Table S2** while those for single-point qualitative analyses appear in ***SI Appendix*, Fig. S5** and **Table S3**. **Table 2** highlights the complete BLI workflow.

**Table 2.** Experimental Design of BLI Binding Studies.

| Sensor | Ligand(s) | Analyte(s) | Quantification | Figure(s) |
| --- | --- | --- | --- | --- |
| NiNTA | cCpE-<sup>His10</sup> | claudin-Biotin | Multi-point | S5A-E |
| NiNTA | CpE-<sup>His10</sup> | claudin-Biotin | Single-point | S5F |
| NiNTA | cCpE-<sup>His10</sup> | COP-1 | Single-point | S5G |
| SA | claudin-Biotin | COP-1 | Single-point | S5H |
| NiNTA | claudin-Biotin/cCpE-<sup>His10</sup> complexes | COP-1 | Single-point | S5I |
| NiNTA | claudin-Biotin/CpE-<sup>His10</sup> complexes | COP-1 | Single-point | S5J |
| NiNTA | claudin-Biotin/cCpE-<sup>His10</sup> complexes | COP-1 | Multi-point | S5K-O |
| SA | hCLDN-4-Biotin/cCpE-<sup>His10</sup> complexes | COP-1 | Multi-point | S5P |

For claudin binding to enterotoxin measurements the time courses for association and dissociation were fit to a 1:1 binding model using the BLItz Pro 1.3 Software. COP-1 binding to claudins and claudin/enterotoxin complexes measurements, however, did not fit a 1:1 binding model, so a 2:1 heterogenous ligand model was used. This model assumes COP-1 binds at two independent antigen sites with different rate constants, which can be explained by independent binding of each COP-1 chain to the complex or two binding modes that rely on a conformational switch in the claudin/cCpE complex to expose the COP-1 binding epitope. Thus, two affinity and rate constants are reported. At the protein concentrations used, no significant non-specific binding of COPs to NiNTA or SA sensors were detected.

### Biochemical Characterization

For biochemical analyses we used samples prepared as described above used for biophysical analyses. Here, 300 μL of 600 nM pre-formed claudin-_Biotin_/cCpE-_His10_ complexes (∼7 μg) were extracted from 1.5 wells of the 96-well plate. These samples, representing five claudin/cCpE complexes, were injected onto a TSKgel QC-PAK GFC 300 SEC column (Tosoh Bioscience) equilibrated in BLI buffer and run for 18 minutes at a flow rate of 0.5 mL/min. Another 300 μL of these same five claudin/cCpE complexes were added to a microcentrifuge tube and 1 molar excess COP1 (∼10 μg) was added. These samples were nutated at 20°C for 1 hour as with BLI to form complexes. After time, these samples, representing five claudin/cCpE/COP-1 complexes, were injected onto the SEC column. As a control, 300 μL of 600 nM pre-formed hCLDN-4-_Biotin_/cCpE-_His10_ complexes was bound to 1 molar excess of COP-2, a cCpE-binding sFab, incubated at 20°C for 1 hour, then injected on SEC. Retention of complexes was assessed by observing decreases to the elution times of claudin/cCpE/sFab complexes compared to uncomplexed claudin/cCpE peak fractions.

## Data Availability

The structural coordinates and cryo-EM maps of the hsCLDN-4/cCpE/COP-1/Nb and hsCLDN-4/cCpE/COP-1 complexes will be deposited in the Protein Data Bank (PDB) and the Electron Microscopy Data Bank (EMDB) before submission to a peer-reviewed journal.

## Acknowledgments

Research reported in this publication was supported by the National Institute of General Medical Sciences of the National Institutes of Health under Award Number R35GM138368 (to A.J.V.) and R01GM117372 (to A.A.K). The content is solely the responsibility of the authors and does not necessarily represent the official views of the National Institutes of Health. Grant support, cryo-EM training, and this material is based upon work supported by the National Science Foundation under Grant OIA-2131902 (to A.J.V.). We are grateful to the University of Chicago Advanced Electron Microscopy Core Facility (RRID:SCR_019198) for providing time and support for cryo-EM data collection. We are also grateful to Craig Yoshioka and Sean Mulligan at the Pacific Northwest Cryo-EM Center (PNCC) for training in cryo-EM acquired via NSF grant OIA-2131902. A portion of this research was supported by NIH grant U24GM129547 and performed at the PNCC at OHSU and accessed through EMSL (grid.436923.9), a DOE Office of Science User Facility sponsored by the Office of Biological and Environmental Research.

## Supplementary Information (SI) Appendix

### SI Appendix, Figures and Tables

**Figure S1.**
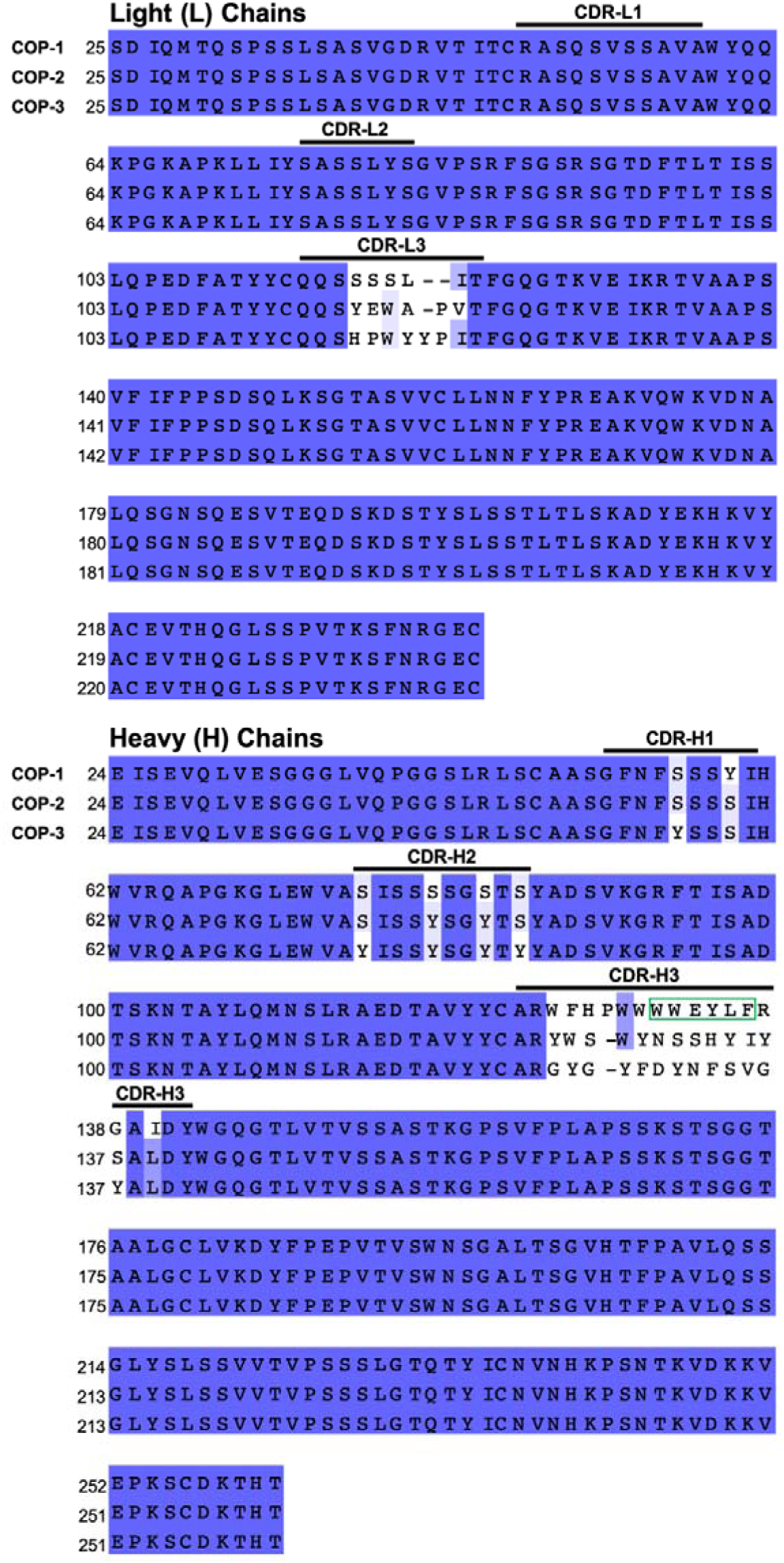
Sequence Alignment of COPs. The L and H chains of COP-1, -2, and -3 were aligned and highlighted within are the CDR-1-3 regions. The amphipathic helix sequence of CDR-H3 in COP-1 in boxed (green).

**Figure S2.**
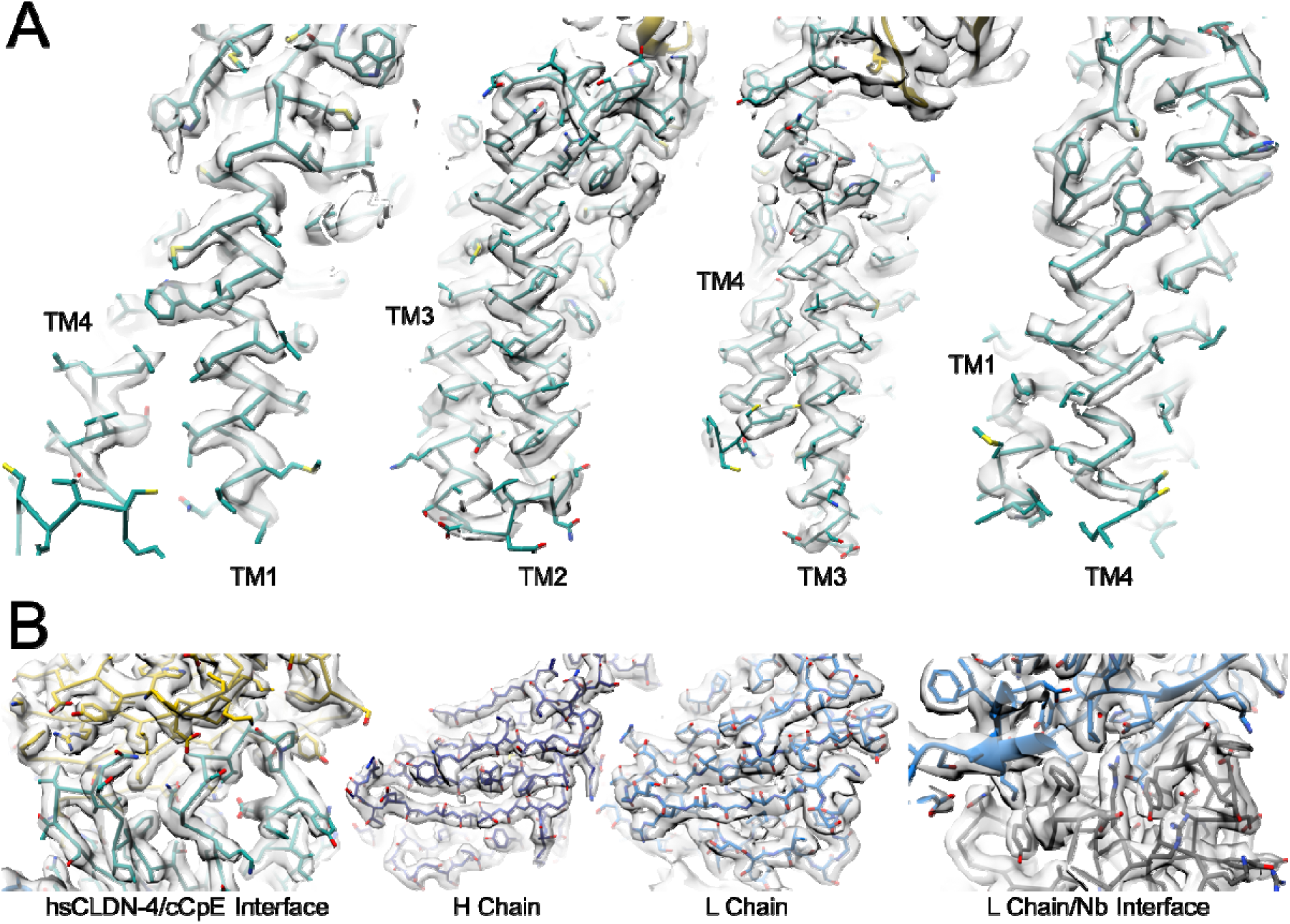
High Resolution Features of hsCLDN-4/cCpE/COP-1/Nb Map. (A) Final cryo-EM map (light grey) of the hsCLDN-4 TM region (teal) within the LMNG micelle. Depicted are the map features around the modeled four individual IZ-helical TMs. (B) Final cryo-EM map of extracellular areas. Depicted are the map features (light grey) around the modeled hsCLDN-4 (teal) and cCpE (yellow) interface, COP-1 H chain (dark blue) and L chain (light blue) -sheets, and the interface between the Nb (grey) and COP-1 L chain (light blue).

**Figure S3.**
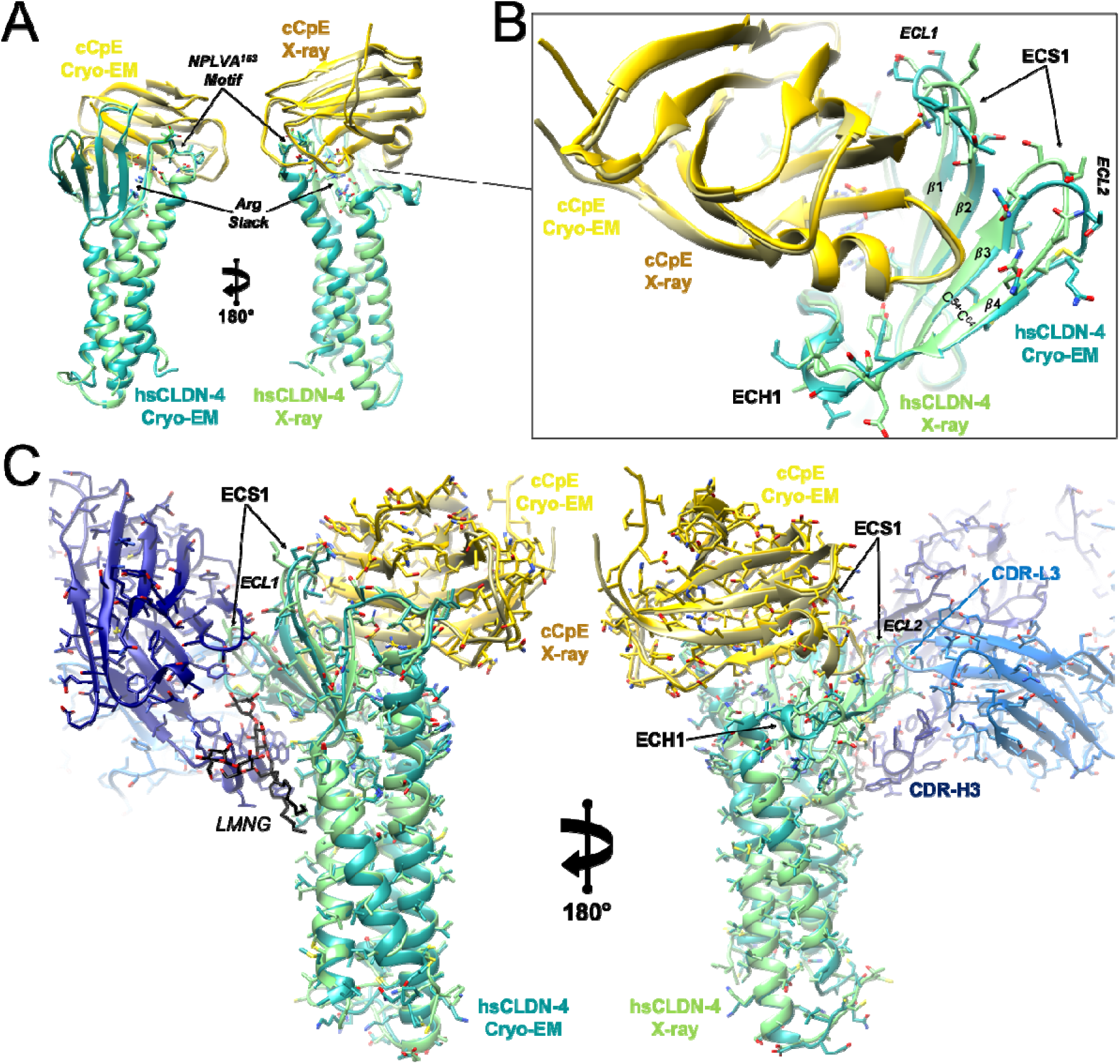
Structural Alignment of the hsCLDN-4/cCpE Complexes from X-ray Crystallography and Cryo-EM. (A) Overall alignment of hsCLDN-4 and cCpE from X-ray (light green/copper) and Cryo-EM (teal/gold). (B) Zoom-in depicting the structural differences between two extracellular loops (ECL) within ECS1 between the X-ray and cryo-EM structures, colored as in A. (C) Structural overlays of the X-ray and cryo-EM structures, colored as in A, with COP-1 (blue) shown for reference.

**Figure S4.**
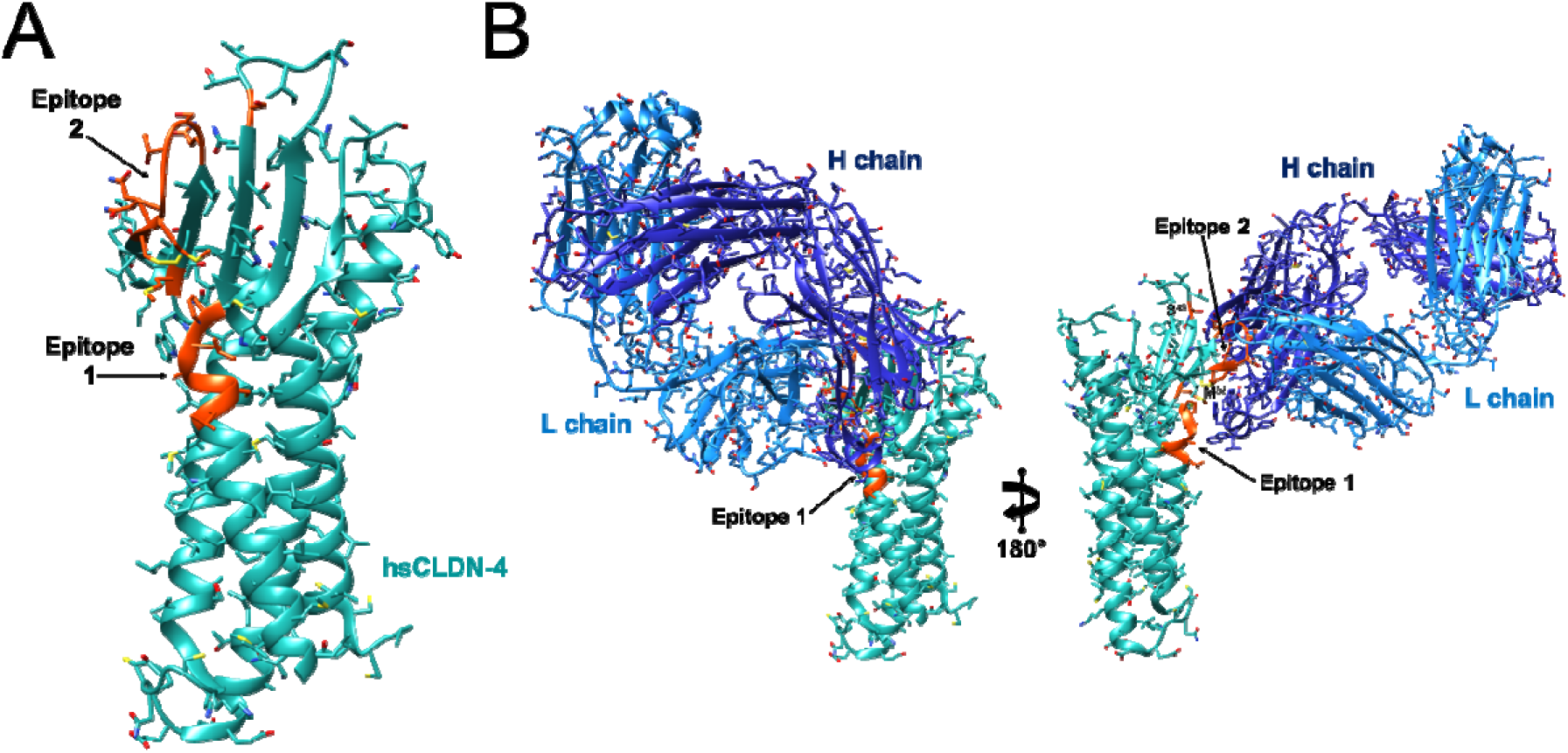
COP-1 Binding Epitopes on hsCLDN-4. (A) The hsCLDN-4 (teal) portion of the structure depicting two epitopes (orange) that interact directly with COP-1. (B) hsCLDN-4 is colored as in A with the COP-1 H (dark blue) or L chains (light blue) interacting areas/residues shown binding epitopes 1 and 2 for reference.

**Figure S5.**
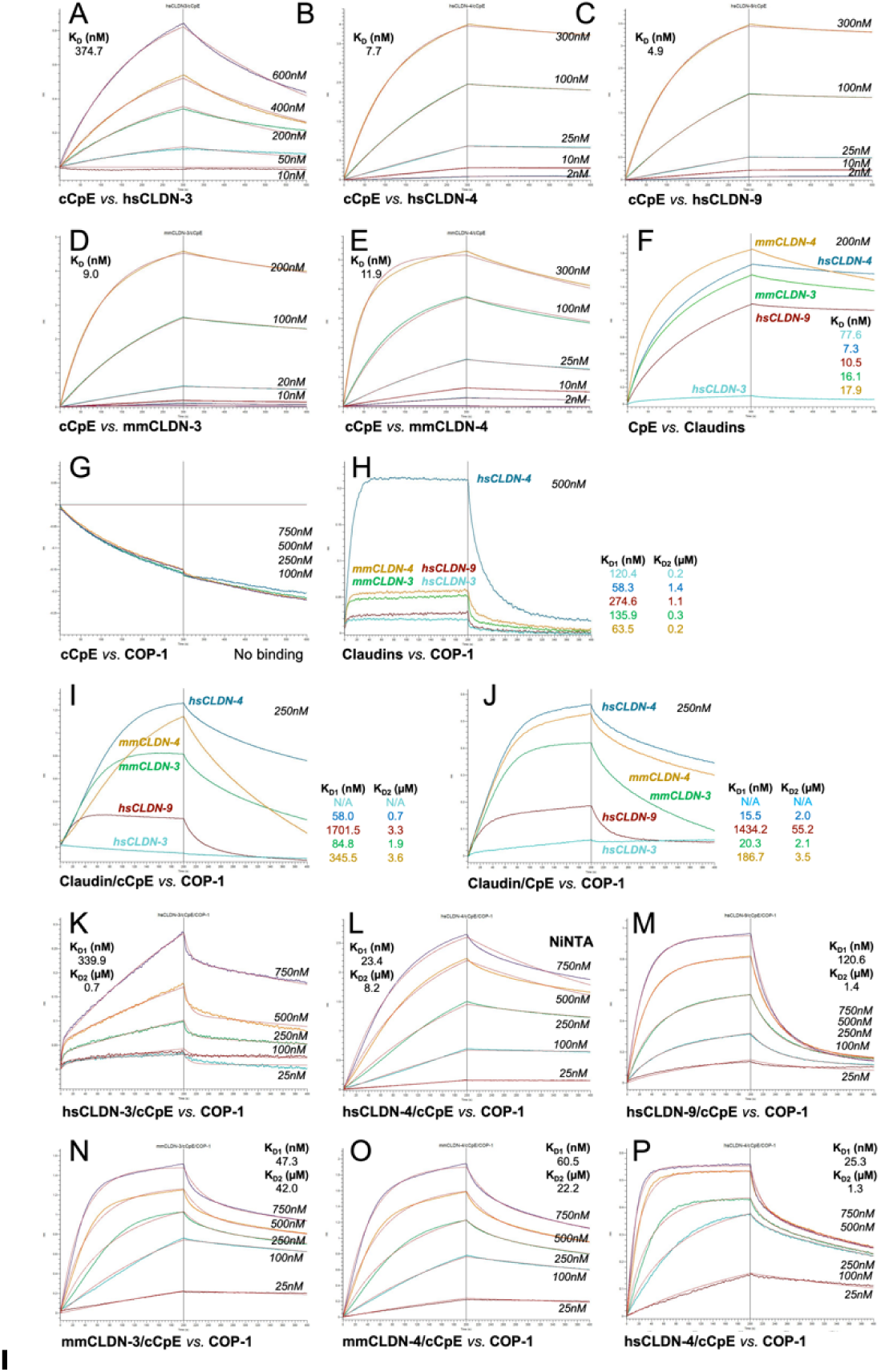
BLI Sensograms Depicting Binding of Various Protein Complexes. Sensograms are grouped accordingly: A-F, Full and semi-quantitative experiments of claudin binding to enterotoxins in the absence of COP-1; G-J, Qualitative binding of COP-1 to cCpE, claudins, and claudin/enterotoxin complexes; and K-P, Quantitative kinetic binding experiments of COP-1 binding to claudin/cCpE complexes. Specifically, cCpE_-His10_ immobilized on NiNTA sensors bound to (A) human claudin-3, (B) human claudin-4, (C) human claudin-9, (D) mouse claudin-3, and (E) mouse claudin-4. (F) shows CpE_-His10_ immobilized on NiNTA sensors binding against 200 nM of aforementioned claudins. Inset table reports the estimated K_D_ from these single-point analyses. All claudin/enterotoxin binding results (A-F) were fit using a 1:1 binding model. (G) shows cCpE_-His10_ immobilized on NiNTA sensors binding against 100-750 nM COP-1 without claudins. (H) shows claudin_-Biotin_ immobilized on SA sensors binding against 500 nM COP-1 without cCpE. Inset table reports the estimated K_D’s_ from these single-point analyses. (I) shows cCpE_-His10_/claudin complexes immobilized on NiNTA sensors binding against 250 nM COP-1. Inset table reports the estimated K_D’s_ from these single-point analyses. (J) shows CpE_-His10_/claudin complexes immobilized on NiNTA sensors binding against 250 nM COP-1 and inset table reports the estimated K_D’s_ from these single-point analyses. cCpE_-His10_/human claudin-3 (K), cCpE_-His10_/human claudin-4 (L), cCpE_-His10_/human claudin-9 (M), cCpE_-His10_/mouse claudin-3 (N), and cCpE_-His10_/mouse claudin-4 (O) complexes immobilized on NiNTA sensors bound to COP-1. (P) shows cCpE_-His10_/human claudin-4_-Biotin_ immobilized on SA sensors bound to COP-1. All COP-1 binding results (H-P) were fit using a 2:1 heterogenous ligand binding model.

**Figure S6.**
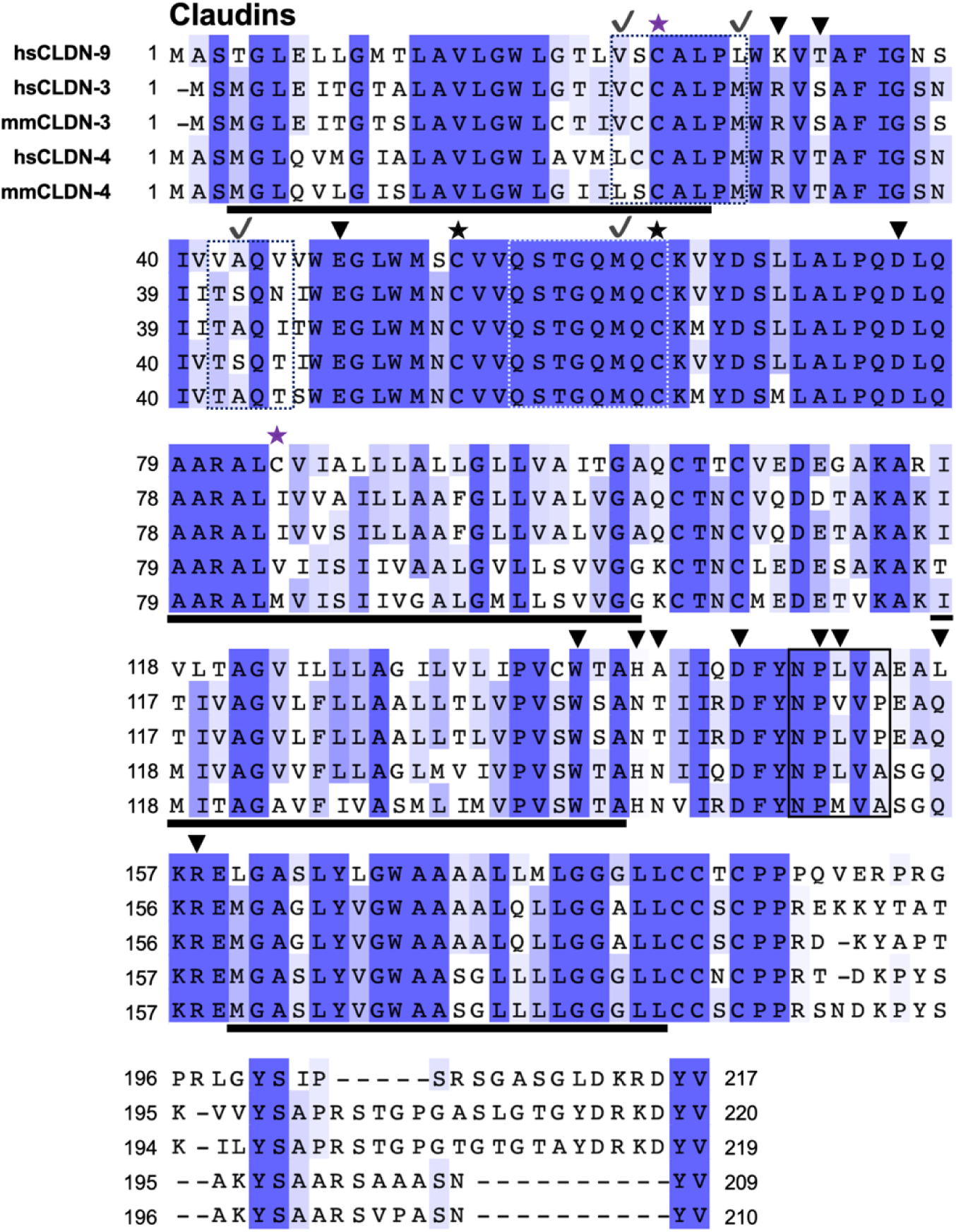
Sequence Alignment of Claudins. The sequences of claudins used in this study were aligned and highlighted within the sequences are the following: Ill (black) represent the conserved disulfide bond in ECS1; Ill (purple) represent the disulfide bond in TM1-TM2 in hsCLDN-9; ▼(black) depict residues that constitute the cCpE-binding motif; box (black) highlights the NPLVA^153^ motif; and (black) shows residues predicted to drive COP-1 interactions with hsCLDN-4. Underlined bars (black) show the residues comprising TM1-4 between all five claudins.

**Figure S7.**
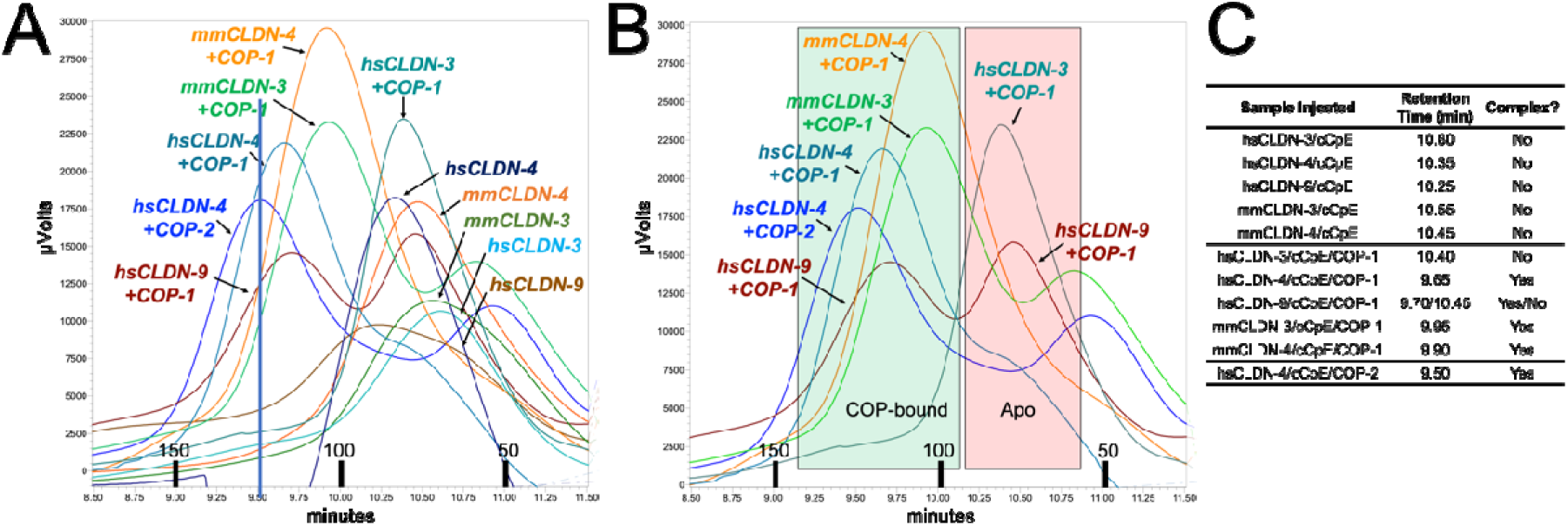
Biochemical Validation of COP-1 Biophysical Findings. Complexes of claudins and cCpE_-His10_ from BLI 96-well plates were injected alone and incubated with excess COP-1, then each sample was injected onto a SEC column equilibrated in BLI buffer. Decreases in peak retention times correlate to increased molecular masses as a result of COP-1 binding to claudin/cCpE complexes. Black bars on the X-axis represent approximate MWs of complexes in kDa. (A) SEC peaks of hsCLDN-3/cCpE (light blue), hsCLDN-4/cCpE (dark blue), hsCLDN-9/cCpE (brown), mmCLDN-3/cCpE (green), and mmCLDN-4/cCpE (orange) from BLI. Overlaid are SEC peaks of hsCLDN-3/cCpE (light teal), hsCLDN-4/cCpE (dark teal), hsCLDN-9/cCpE (maroon), mmCLDN-3/cCpE (light green), and mmCLDN-4/cCpE (light orange) from BLI incubated with COP-1. The control sample of hsCLDN-4/cCpE bound to COP-2 (blue) is shown to validate that the peaks eluting <10 minutes represent sFab-bound complexes. (B) Data from just incubations with excess COPs are shown and colored as in A, and boxes representing COP-bound (green) or unbound (red) are added for clarity. (C) Table inset reporting peak retention times of various samples.

**Figure S8.**
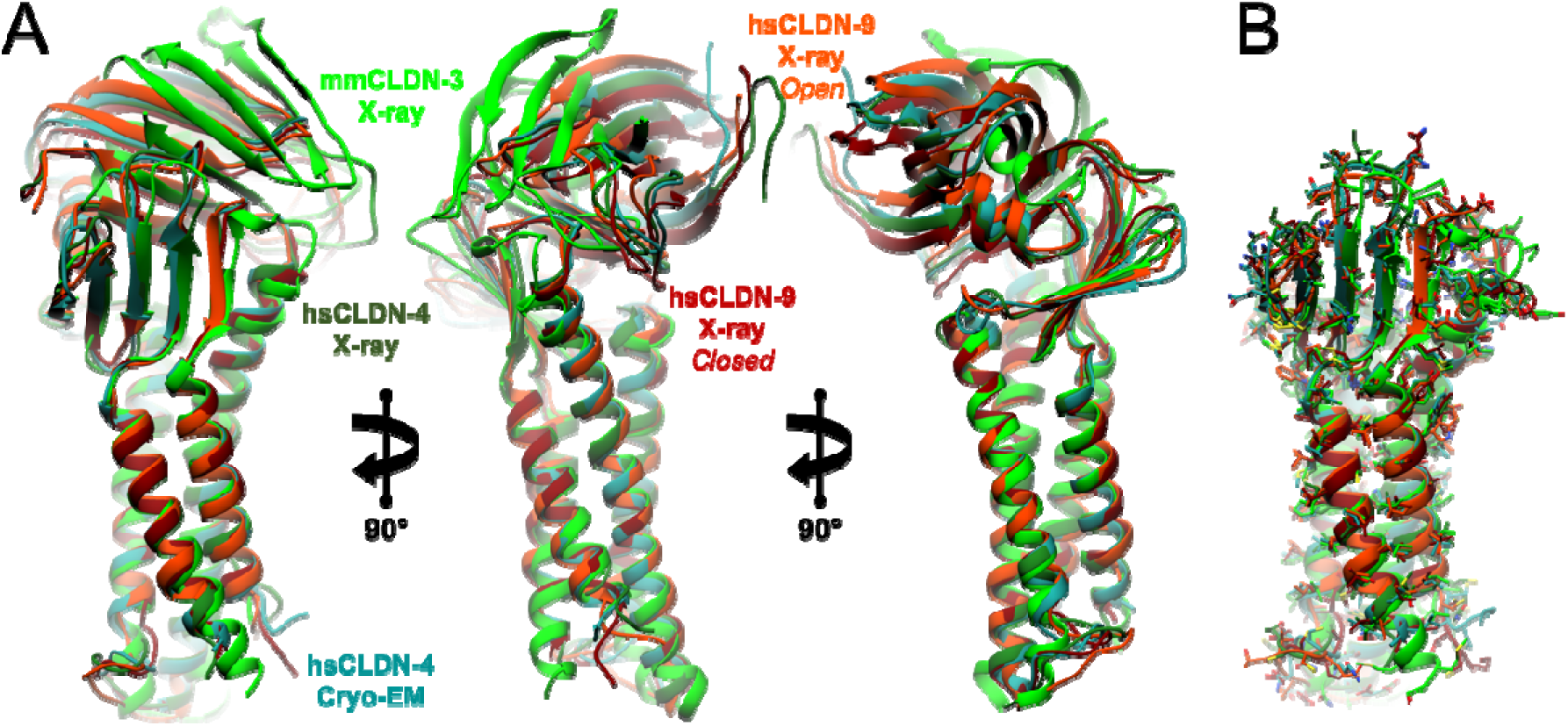
Conformation Comparison between Claudin/cCpE Complexes. Structures of claudins in complex with cCpE as determined by X-ray crystallography and cryo-EM were compared. (A) hsCLDN-4/cCpE (teal) from cryo-EM; and PDB ID: 7KP4 hsCLDN-4/cCpE (green), PDB ID: 6AKE mmCLDN-3/cCpE (light green), and PDB ID: 6OV2 and 6OV3 hsCLDN-9/cCpE (red and orange) from X-ray crystallography are shown and rotated 180°. (B) For clarity, cCpE is removed from all five structures, revealing minimal alterations in the global structure of claudins.

**Figure S9.**
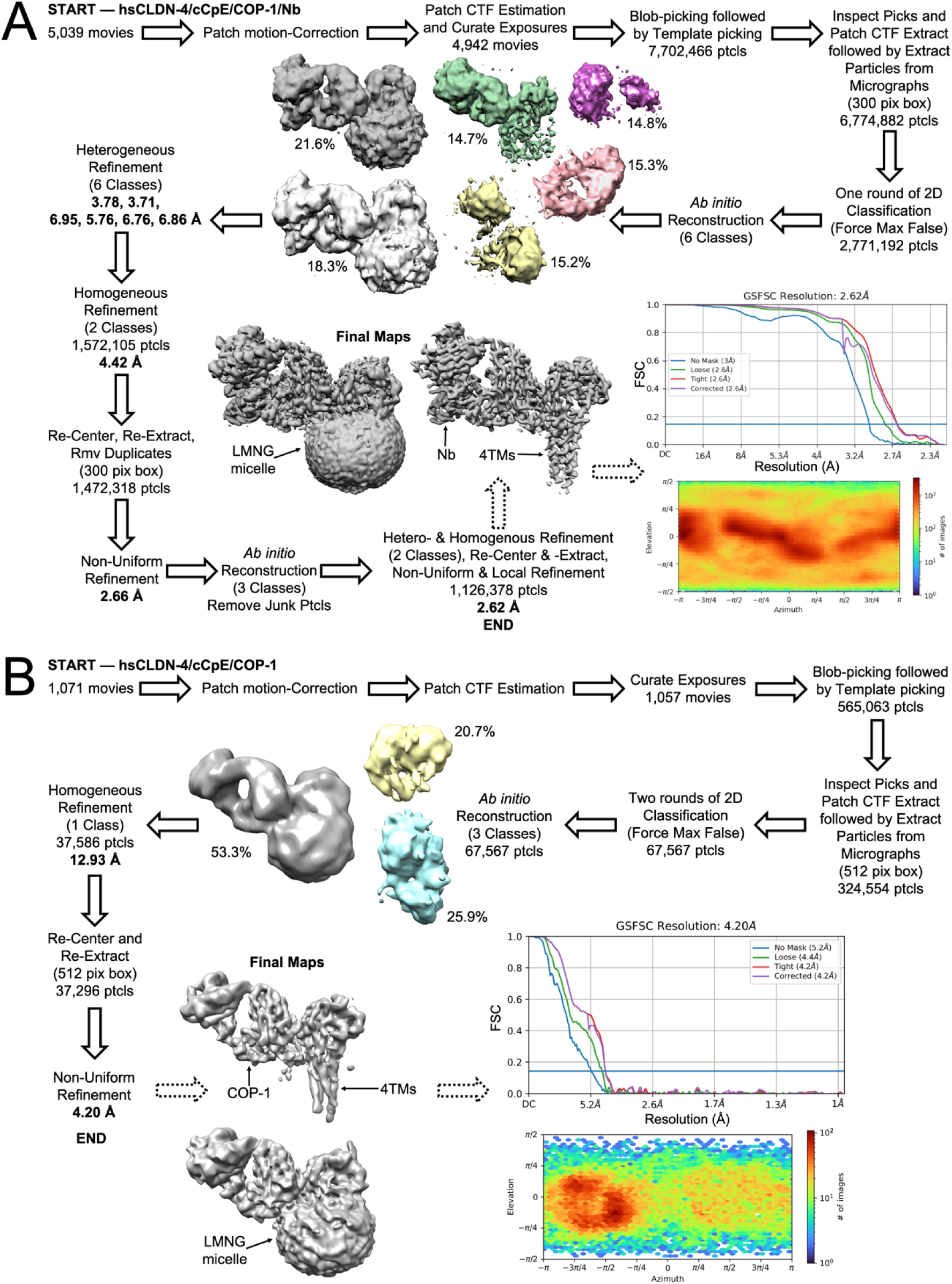
Cryo-EM Data Processing Workflows for hsCLDN-4/cCpE/COP-1 Complexes. (A) Workflow for the hsCLDN-4/cCpE/COP-1/Nb complex. (B) Workflow for the hsCLDN-4/cCpE/COP-1 complex. Fourier Shell Correlation (FSC) curves from gold-standard refinement of both complexes are shown with the 0.143 FSC cutoff indicated by line (blue). Plots of the angular distribution of particles in the final refinement are shown below FSC curves.

**Table S1.**
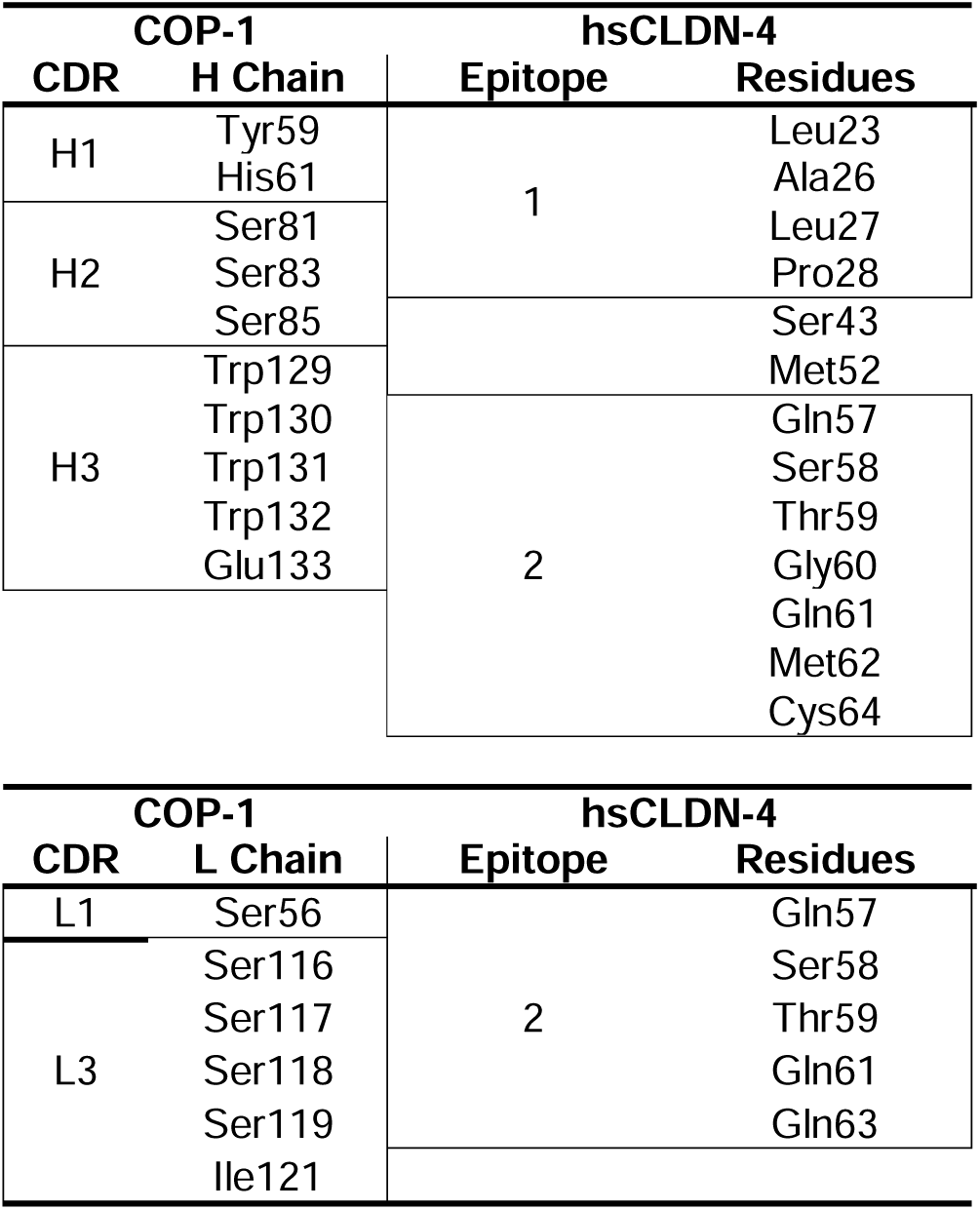
Side Chains of Significance in hsCLDN-4 and COP-1 Interactions. Side chains that interact according to structural analyses are shown below. An interaction is defined as a polar or non-polar interaction that occurs between two atoms in these residues at a distance range between 1.4-4.0 Å.

**Table S2.** BLI Binding Results For Multi-Point Analyses with Kinetics.

| Table 1 and Fig. S5A-E Data |  |  |  |  |
| --- | --- | --- | --- | --- |
| Complex | k<sub>on</sub><br>(1/Ms) | k<sub>off</sub> (1/s) | K<sub>D</sub> (nM) | t<sub>1/2</sub><br>(min) |
| hsCLDN-3<br>/ cCpE | 6.0×10<sup>3</sup> | 2.3×10<sup>-3</sup> | 374.7 ± 2.1 | 5.0 |
| hsCLDN-4<br>/ cCpE | 2.7×10<sup>4</sup> | 2.1×10<sup>-4</sup> | 7.7 ± 0.1 | 55.0 |
| hsCLDN-9<br>/ cCpE | 2.7×10<sup>4</sup> | 1.3×10<sup>-4</sup> | 4.9 ± 0.1 | 88.9 |
| mmCLDN-3<br>/ cCpE | 5.0×10<sup>4</sup> | 4.5×10<sup>-4</sup> | 9.0 ± 0.1 | 25.7 |
| mmCLDN-4<br>/ cCpE | 6.9×10<sup>4</sup> | 8.2×10<sup>-4</sup> | 11.9 ± 0.1 | 14.1 |

| Table 1 and Fig. S5K-P Data |  |  |  |  |  |  |  |
| --- | --- | --- | --- | --- | --- | --- | --- |
| Complex | k<sub>on 1</sub><br>(1/Ms) | k<sub>off 1</sub><br>(1/s) | K<sub>D 1</sub> (nM) | k<sub>on 2</sub><br>(1/Ms) | k<sub>off 2</sub><br>(1/s) | K<sub>D 2</sub> (μM) | t<sub>1/2</sub><br>(min) |
| hsCLDN-3•cCpE /<br>COP-1 | 2.2×10<sup>5</sup> | 7.9×10<sup>-2</sup> | 339.9 ± 9.4 | 7.6×10<sup>4</sup> | 6.1×10<sup>-2</sup> | 0.7 ± 1.5 | 0.1 |
| hsCLDN-4•cCpE /<br>COP-1 | 2.4×10<sup>4</sup> | 5.5×10<sup>-4</sup> | 23.4 ± 1.7 | 2.6×10<sup>3</sup> | 2.1×10<sup>-2</sup> | 8.2 ± 6.6 | 21.2 |
| hsCLDN-9•cCpE /<br>COP-1 | 3.6×10<sup>4</sup> | 4.9×10<sup>-3</sup> | 120.6 ± 0.8 | 2.3×10<sup>4</sup> | 3.3×10<sup>-2</sup> | 1.4 ± 0.1 | 2.4 |
| mmCLDN-3•cCpE /<br>COP-1 | 2.7×10<sup>4</sup> | 1.1×10<sup>-3</sup> | 47.3 ± 1.0 | 0.7×10<sup>3</sup> | 2.8×10<sup>-2</sup> | 42.0 ± 24.9 | 10.6 |
| mmCLDN-4•cCpE /<br>COP-1 | 1.9×10<sup>4</sup> | 9.7×10<sup>-7</sup> | 60.5 ± 2.1 | 1.0×10<sup>3</sup> | 1.9×10<sup>-2</sup> | 22.2 ± 4.0 | 11.9 |
| hsCLDN-4•cCpE /<br>COP-1<br>(SA) | 8.3×10<sup>4</sup> | 2.1×10<sup>-3</sup> | 25.3 ± 0.2 | 4.0×10<sup>4</sup> | 5.3×10<sup>-2</sup> | 1.3 ± 0.1 | 5.5 |

**Table S3.**
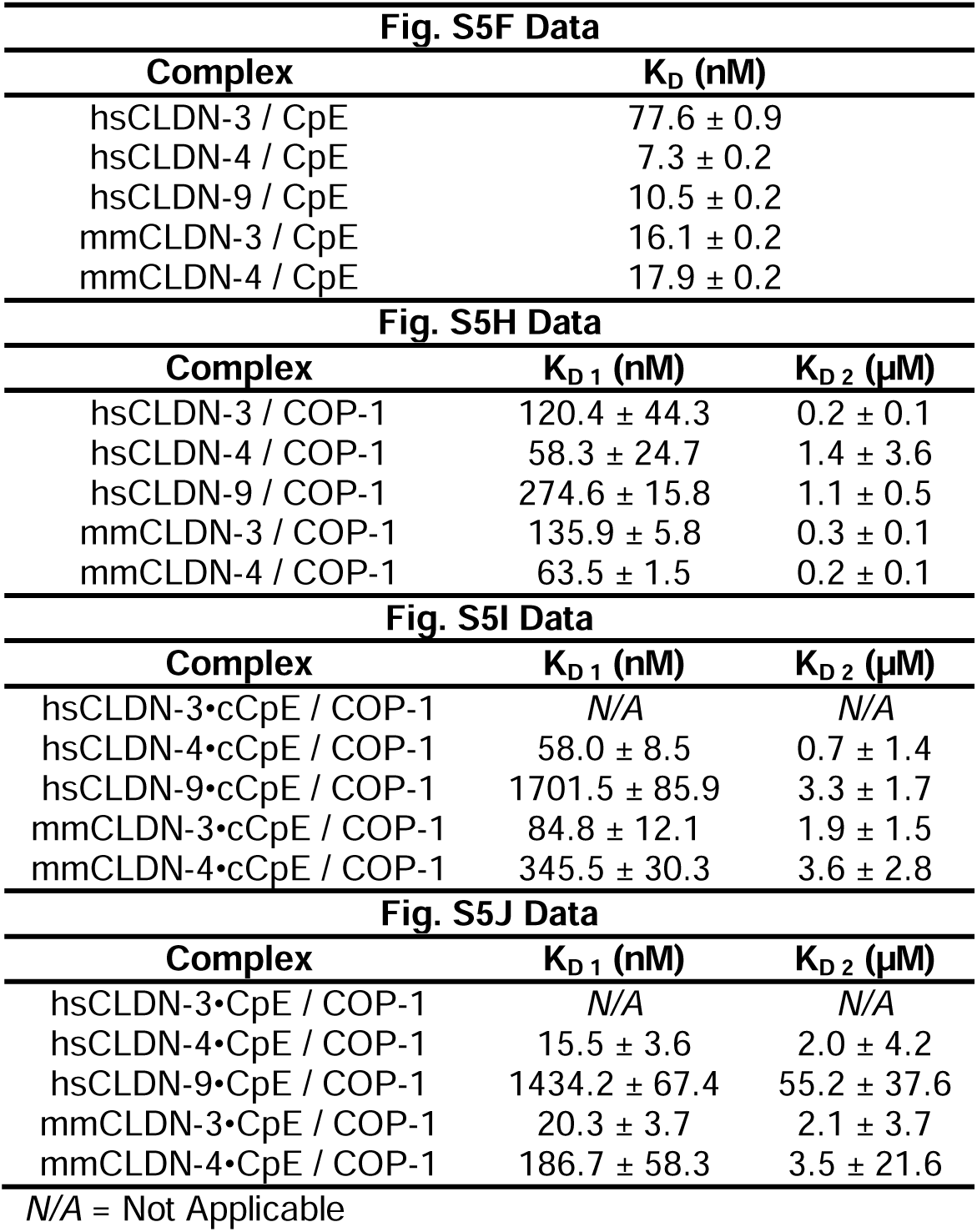
BLI Binding Results For Single-Point Analyses.

**Table S4.** Cryo-EM Data Collection, Refinement and Validation Statistics.

|  | hsCLDN-4/cCpE/COP-1/Nb | hsCLDN-4/cCpE/COP-1 |
| --- | --- | --- |
| **Data collection and processing** |  |  |
| Magnification | 81,000 | 92,000 |
| Voltage (kV) | 300 | 300 |
| Electron exposure (e<sup>-</sup>/Å<sup>2</sup>) | 60 | 60 |
| Defocus range (μm) | 0.9 - 2.1 | 0.8 - 2.2 |
| Pixel size (Å) | 1.068 | 0.507 |
| Symmetry imposed | C1 | C1 |
| Number of micrographs | 5,039 | 1,071 |
| Initial particle images (no.) | 7,702,466 | 565,063 |
| Final particle images (no.) | 1,126,378 | 37,296 |
| Map resolution (Å) | 2.62 | 4.20 |
| FSC threshold | 0.143 | 0.143 |
| **Refinement** |  |  |
| Model resolution (Å) | 2.62 | 4.15 |
| FSC threshold | 0.143 | 0.143 |
| Model composition |  |  |
| Non-hydrogen atoms | 6647 | 5721 |
| Protein residues | 864 | 745 |
| B factors (Å<sup>2</sup>) |  |  |
| Protein | 123.84 | 238.34 |
| Ligand | 158.33 | 270.75 |
| Water | 109.85 | N/A |
| R.M.S. deviations |  |  |
| Bond lengths (Å) | 0.007 | 0.008 |
| Bond angles (°) | 1.124 | 1.144 |
| Validation |  |  |
| MolProbity score | 1.94 | 1.84 |
| Clashscore | 7.74 | 11.58 |
| Poor rotamers (%) | 3.04 | 0.00 |
| Ramachandran plot |  |  |
| Favored (%) | 97.19 | 96.20 |
| Allowed (%) | 2.81 | 3.80 |
| Disallowed (%) | 0.00 | 0.00 |

N/A = Not Applicable

